# Rethink the Sink: Urban cores act as leaky sinks to maintain regional gene flow in coyotes

**DOI:** 10.64898/2026.09.24.753963

**Authors:** Summer E. Vance, Niamh Quinn, Paul Stapp, Bridgett vonHoldt, Christopher J Schell

## Abstract

Urbanization is an accelerating evolutionary force, yet the patterns and drivers of urban gene flow remain largely obscured. This leaves a critical gap in testing the interplay between demographic and genetic models of urban connectivity. Specifically: the urban fragmentation model, where structural barriers restrict gene flow and drive genetic drift; the urban facilitation model, where permeable matrices enhance functional connectivity; and the urban sink hypothesis, a demographic model where high mortality creates unsustainable population sinks. We used high-resolution RAD-capture data to conduct landscape genomics and gene flow analyses to resolve these models and investigate contemporary evolution in urban coyotes (*Canis latrans*) across the greater Los Angeles, CA, USA metropolitan area. By incorporating fine-scale environmental, social-ecological, and historical variables of the built environment, we rigorously assessed how urban heterogeneity shapes intra-city evolutionary dynamics. We found weak population structure, heavily influenced by family groups. There was extensive landscape permeability and gene flow across the study area including across four major highways. We also identified heavily asymmetric gene flow from exurban source populations into the urban core, supporting the urban sink paradigm. However, rather than acting as a genetic dead-end, the urban matrix showed striking functional heterogeneity: one urban core functioned as a "genetic sponge," absorbing unique transient alleles from multiple surrounding sources, and both urban cores operated as permeable “leaky sinks,” exporting critical gene flow to other populations. Consequently, urban demographic sinks can simultaneously act as vital genetic bridges. Our findings reject a pure urban fragmentation model while demonstrating that demographic sink dynamics and genetic facilitation are not mutually exclusive. This demonstrates that high-turnover urban sinks can buffer regional biodiversity against genetic erosion, highlighting the need to prioritize matrix permeability and inter-patch connectivity in urban conservation strategies.

**Significance:** Urban landscapes are widely assumed to act either as impenetrable barriers that accelerate genetic isolation or as demographic "sinks" that trap dispersing wildlife. Studying coyotes across Greater Los Angeles, we demonstrate that demographic sink dynamics and functional genetic connectivity are not mutually exclusive. Instead, high-turnover urban matrices exhibited marked functional heterogeneity:urban cores can absorb and harbor transient genetic diversity from surrounding regions while simultaneously operating as "leaky sinks" that export outward gene flow. By revealing that urban demographic sinks can function as critical genetic bridges, this work transforms our understanding of urban evolutionary dynamics and emphasizes matrix permeability as a vital priority for regional biodiversity conservation.

**Graphical abstract:** 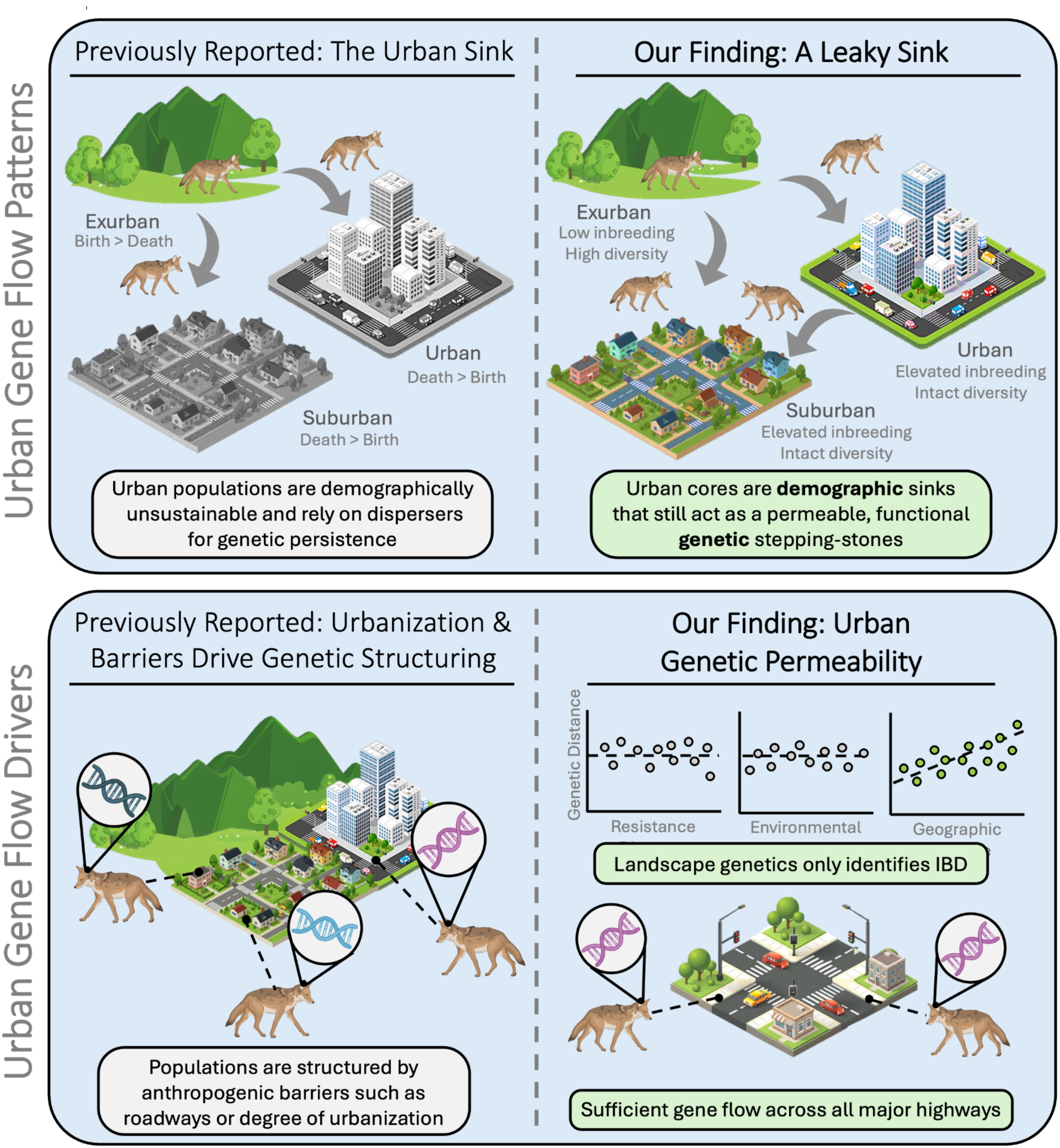

## Introduction

Urbanization is rapidly reshaping the distribution of biodiversity as well as the evolutionary trajectories of wildlife globally (1–3). At the core of urban evolutionary dynamics is gene flow, which governs how genetic variation is partitioned, maintained, or eroded across landscapes. Because gene flow is driven by dispersal and subsequent survival, the structural architecture and social-ecological matrix of cities dictate population connectivity by altering dispersal and demographics (4–6). Consequently, evaluating urban connectivity requires untangling the interplay between genetic models of landscape permeability and demographic models of population persistence. In mammals, urbanization is often associated with decreased genetic diversity and thus, eroded evolutionary adaptive capacity (7–9). The urban fragmentation model assumes this pattern is driven by anthropogenic barriers that restrict gene flow into and within urban environments, accelerating genetic drift and loss of genetic diversity (4). Alternatively, the urban facilitation model states the opposite: urbanization acts as a conduit to increase dispersal rates and resulting gene flow. At the demographic level, the urban sink models posit that human-dominated landscapes act as sinks, where wildlife populations experience high mortality and are suboptimally maintained only through continuous immigration from nearby exurban and/or rural source populations (10–12). Resolving how these landscape-movement interactions shape gene flow is critical. Many cities are situated within global biodiversity hotspots and are increasingly recognized as critical nodes for regional conservation (13,14). If urban matrices act primarily as impenetrable barriers (fragmentation), conservation must prioritize restoring intra-city structural corridors to prevent genetic erosion. If they augment dispersal (facilitation), urban networks can independently sustain regional gene flow. However, if cities operate as high-turnover demographic traps (urban sinks), the long-term genetic persistence of urban biodiversity remains fundamentally tied to preserving surrounding exurban source populations.

The structural and social-ecological heterogeneity of cities is known to alter animal movement (15) and contribute to high incidences of human-induced death (16,17). However, both the overarching patterns and specific drivers of gene flow remain poorly understood. Teasing apart the demographic processes that drive genetic patterns requires a layered analytical framework including high-resolution spatial, genomic, and social-ecological data. Prevailing literature primarily relies on a simplified urban-rural dichotomy that masks the immense heterogeneity of cities (18). While studies have detected population structure in intra-urban carnivores, these analyses have largely relied on coarse environmental proxies or focused on isolated linear barriers (e.g., highways). This reductionist approach overlooks both the inherent environmental complexity of cities, as well as the tight coupling of evolution and social-ecological phenomena (19). Social-ecological drivers such as infrastructure, population density, pollution burden, and historical policies (e.g. redlining) have demonstrable effects on contemporary biodiversity, occupancy, and movement patterns (15,20–22), yet are rarely incorporated into urban evolutionary studies (18, but see 19). Furthermore, because cities have developed on an exceedingly short evolutionary timescale, failing to integrate historical landscape data limits our ability to distinguish between the effects of past and present urbanization on observed genetic patterns (25).

To resolve dynamics of contemporary gene flow and urban evolution, we leverage the Greater Los Angeles metropolitan area (hereafter referred to as Greater LA), a megacity in California, USA, offering a high-contrast gradient between high-density urban cores and adjacent wildlands. As the archetypal and widespread urban adapter, the coyote (*Canis latrans*) serves as an our focal species. Coyotes are uniquely suited for landscape genomic investigations because they combine high dispersal capacity and generalist ecology with an exceptional ability to exploit anthropogenic subsidies and navigate dense urban infrastructure. Consequently, their movement patterns and genetic architecture provide a unique ecological sensor for how modern infrastructure fragments, channels, or facilitates contemporary mammalian gene flow.

Here, we used a multi-scaled population genomic framework to achieve three primary objectives: 1. Assess the baseline distribution of genetic diversity and subpopulation structure across Greater LA; 2. Employ landscape genomics using six fine-scale environmental metrics to identify specific drivers of genetic differentiation; and 3. Model contemporary, directional gene flow to evaluate the genetic permeability of suspected anthropogenic barriers and investigate urban source-sink dynamics. By contrasting these genomic and demographic analyses, we tested the competing predictions of the urban fragmentation, urban facilitation, and urban sink models. If urbanization acts primarily as a barrier (fragmentation model), we hypothesized that coyotes would exhibit distinct, isolated subpopulations bounded by infrastructure, alongside localized signatures of genetic decay and elevated inbreeding. Under this scenario, we predicted that genetic differentiation would be driven by isolation by resistance (IBR) and/or isolation by environment (IBE), allowing us to determine whether structural features (e.g., roads, impervious surfaces) or social-ecological gradients (e.g., pollution, human population density) specifically restrict gene flow beyond simple isolation by distance (IBD). Alternatively, if the urban matrix functions as a permeable conduit (facilitation model), we expected high, multi-directional gene flow across built environments regardless of structural density. Finally, if the city functions primarily as an attractive demographic trap (urban sink hypothesis), we hypothesized that urban core habitat would exhibit weak population structure sustained by continuous, asymmetric immigration from adjacent exurban sources. Ultimately, this work aims to clarify how the physical and social-ecological architecture of megacities shapes the evolutionary trajectories of urban wildlife.

## Results

### Kinship filtering reveals weak genetic structure and high admixture

Inclusion of all individuals (N = 220) resulted in five distinct genetic clusters (Fig S2, Table S1), however, after removing first-degree relatives, population structure was optimally represented by three clusters (K = 3; N = 153). We concluded that the five initial clusters reflected family groups rather than true subpopulations. Consequently, we conducted all downstream analyses using the K = 3 populations, hereafter referred to as North, South, and Central (Fig 1). Individuals exhibited high levels of admixture; nearly half of the unrelated samples (45.10%, N = 69) were classified as admixed, defined as <70% ancestry assigned to any single cluster. Admixed individuals were ubiquitous, occurring in all populations but most concentrated in geographic transition zones between populations and at the periphery of our sampling range (Fig S2).

**Figure 1.**
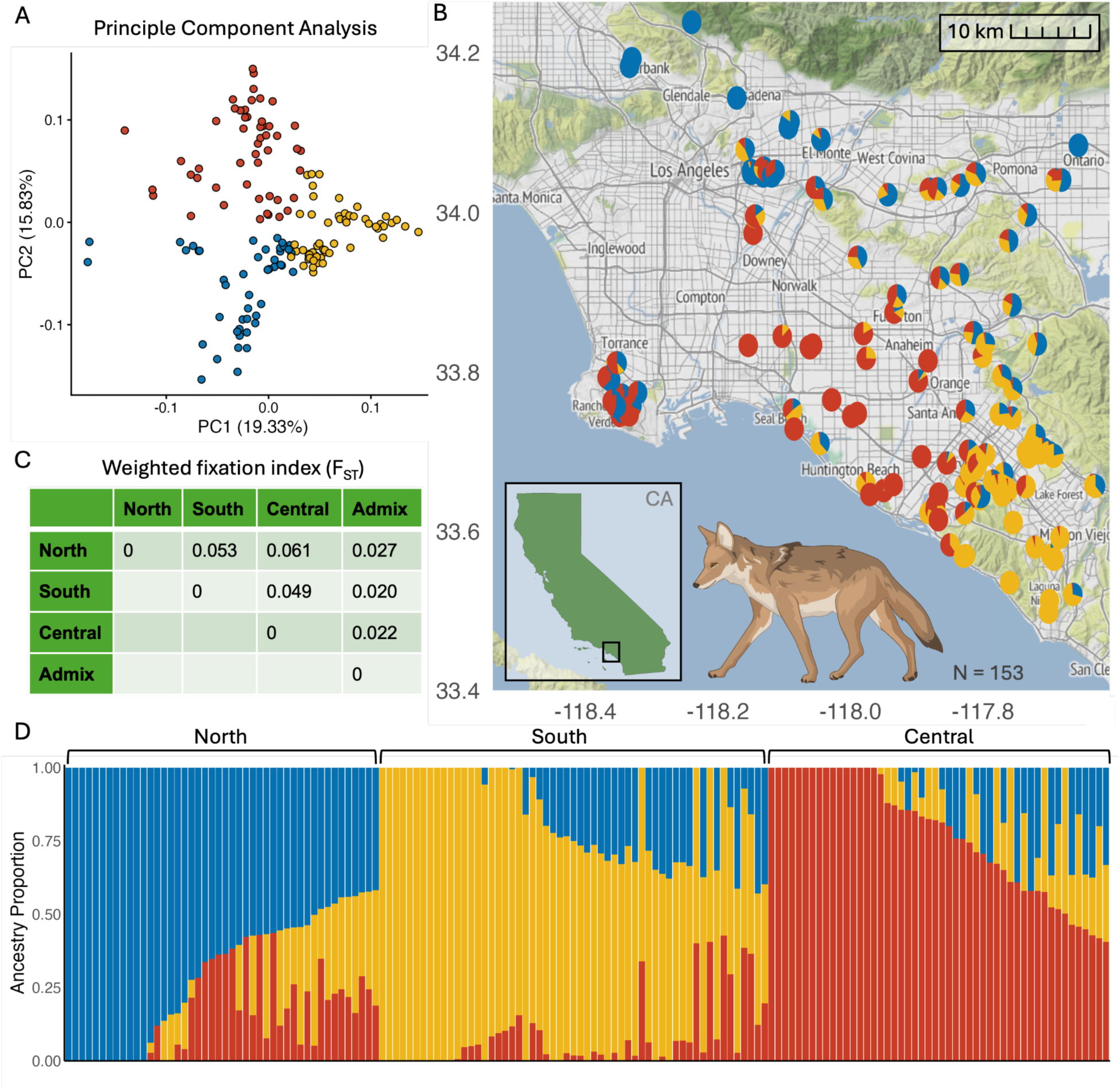
Population genetic structure and spatial admixture. **(A)** Principal coordinate analysis (PCoA) showing genetic differentiation along PC1 (19.33% variance) and PC2 (15.83% variance). Individual points are color-coded by assigned genetic cluster: North (blue), Central (yellow), and South (red). **(B)** Spatial distribution and individual ancestry proportions mapped across the study area. Pie charts represent sampling locations and individual ancestry coefficients derived from K = 3 clustering. Inset map indicates the regional study site within California. **(C)** Pairwise F_ST_ matrix showing weak genetic differentiation among genetic clusters and the admixed group. **(D)** Ancestry proportion bar plot from admixture analysis (K = 3), where each vertical bar represents an individual and colors correspond to cluster assignment probabilities (blue = North, yellow = Central, red = South).

Genetic differentiation among these clusters was weak (Fst range: 0.049–0.061; Table 1). Principal coordinate analysis (PCA) confirmed K=3 clustering, with admixed individuals occupying the intermediate space between populations (Figure 3; Figure S2). The first two axes explained 18.72% and 15.43% of the total genetic variance, respectively. Diversity metrics revealed consistent, significant heterozygote deficits (He > Ho) across all populations (Table 1). Inbreeding levels (Fis) were mild, ranging from 0.05-0.11. Notably, we identified thirty individuals with Fis values >=0.15 and five of these had values >=0.25 (File S1). Allelic richness was consistent across all populations (1.70-1.76). However, normalized private allele counts varied considerably across groups; the Central population exhibited six times fewer private alleles per individual than the South population and nearly thirty-fold fewer than North.

**Table 1.** Population genetics for K = 3. Values represent population estimates derived from K = 3 genetic clusters plus admixed individuals (N = 153 total unrelated samples). Displayed metrics include sample size (N), expected heterozygosity (H_E_), observed heterozygosity (H_O_), results of Welch two-sample t-test statistics comparing He and Ho, inbreeding coefficient (F_IS_), allelic richness, per-individual private alleles, and total private alleles per population.

| Pop | N | H<sub>E</sub> | H<sub>O</sub> | t, df, p* | F<sub>IS</sub> | Allelic richness** | Private alleles** | Total private alleles |
| --- | --- | --- | --- | --- | --- | --- | --- | --- |
| North | 20 | 0.208 ± 0.001 | 0.199 ± 0.0165 | -2.63, 19, **0.017** | 0.0464 ± 0.0788 | 1.760 | 14.80 | 296 |
| South | 32 | 0.208 ± 0.001 | 0.198 ± 0.0182 | -3.29, 31, **0.002** | 0.0493 ± 0.0849 | 1.758 | 3.219 | 103 |
| Central | 32 | 0.208 ± 0.001 | 0.186 ± 0.0215 | -5.87, 31, **1.78 e-6** | 0.105 ± 0.102 | 1.699 | 0.594 | 19 |
| Admix | 69 | 0.208 ± 0.002 | 0.204 ± 0.0137 | -2.40, 68, **0.019** | 0.0188 ± 0.0647 | 1.821 | 23.14 | 1597 |

\*p-values are from a Welch two-sample t-test of unequal variance between H<sub>E</sub> and H<sub>O</sub>
\*\*values are calculated on a per-individual basis to correct for uneven population sizes

**Table 2.**
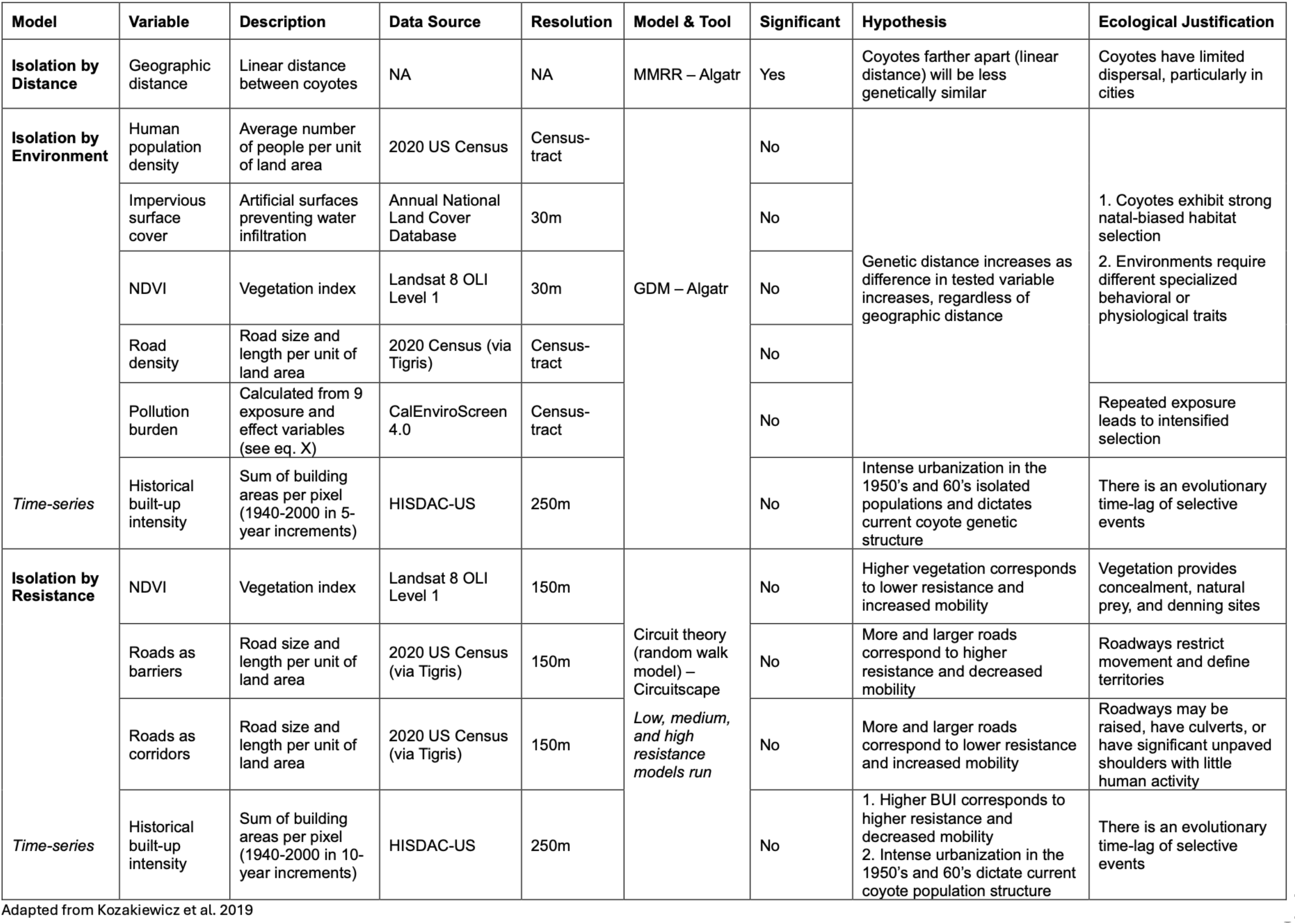
Landscape genetics models and results. Evaluations of Isolation by Distance (IBD), Isolation by Environment (IBE), and Isolation by Resistance (IBR) using Multiple Matrix Regression with Randomizations (MMRR), Generalized Dissimilarity Modeling (GDM) via the R package *Algatr*, and Circuit Theory via *Circuitscape*. Geographic distance was the sole statistically significant driver of genetic differentiation across the landscape.

### Populations experience differing levels of anthropogenic exposure

Environmental exposure across our metrics varied significantly among genetic clusters (Figure S5; Table S1). The North and Central clusters consistently occupied the most heavily urbanized landscapes. The North cluster experienced the highest mean values across all built-environment variables, including human population density (53.84 +/- 13.20), impervious surface cover (61.21 +/- 12.38), pollution burden (46.52 +/- 11.52), and road density (69.95 +/- 11.15). The Central cluster followed closely behind, exhibiting similar urban pressure (48.74 +/- 32.93 human pop density; 58.53 +/- 22.35 impervious surface; 63.05 +/- 22.62 road density).

Conversely, the South cluster occupied substantially less urbanized landscapes, characterized by the lowest pollution burden (28.88 +/- 13.77), lowest impervious surface cover (40.06 +/- 17.58), and the highest vegetation greenness (61.74 +/- 12.43). These findings confirm a trade-off between built environments and vegetative cover. Environmental heterogeneity was highest for human population density (CV ≈ 62%) and lowest for impervious surface cover and NDVI (CV ≈ 37; Table S4). Among groups, North coyotes experienced the most homogeneous environmental conditions, while the South and Central groups faced higher variability (Table S3).

### Isolation by distance is the only significant driver of genetic differentiation

We found that genetic differentiation in greater Los Angeles coyotes is primarily driven by geographic distance (*t* = 49.89, *p* ≤ 0.01, *n_perm_* _=_ 99), with no detectable signal of urban-mediated resistance or environmental sorting across six high-resolution environmental variables. Extended analyses of historical urbanization patterns (1900–2015) revealed no significant signature of historical IBE or IBR (Supplementary Text).

As an initial test of the landscape’s effect on coyote genetics, we calculated correlation coefficients (rho) between genetic statistics and covariate exposure and found that at both the population and individual level, urban covariates and genetic metrics had no significant correlation (Table S6). Multiple matrix regression and generalized dissimilarity models (GDM) confirmed that geographic distance is the dominant driver of genetic dissimilarity, explaining 71–74% of the total model variance (Tables S7-22). In contrast, urban covariates offered minimal explanatory power (11.62–12.57% of total variance). Variable permutation further highlighted the dominance of IBD. Among environmental predictors, road density and impervious surface cover showed the strongest, albeit statistically not significant, contributions to the model (0.12), followed by pollution burden (0.09). NDVI and human population density did not predict any genetic dissimilarity (0.0).

We tested for isolation by resistance (IBR) by comparing pairwise resistance values derived from NDVI, road density (roads as corridors and barriers for movement), and historical built-up intensity (Tables S23 & S24). After calculating IBR while correcting for isolation by distance (IBD) with a partial Mantel test, no IBR model maintained statistical significance, indicating that landscape resistance, as measured by these metrics, does not constrain gene flow.

### Mapping relatedness reveals a highly permeable landscape

We mapped pairwise genetic relatedness across the landscape, categorizing pairs into first- (N = 128), second- (N = 256), and third-degree (N = 861) relatives (Fig 2). Individuals more distantly related than third-degree were classified as unrelated. While average distance between individuals increased with decreasing kinship, there were several major outliers particularly in first- and second-degree relatives, with related individuals occurring over 50km apart across dense urban sprawl (Fig 2). Notably, geographic distance between first-degree relatives serves as a proxy for actual dispersal events, representing the movement of offspring or siblings from their place of birth.

**Figure 2.**
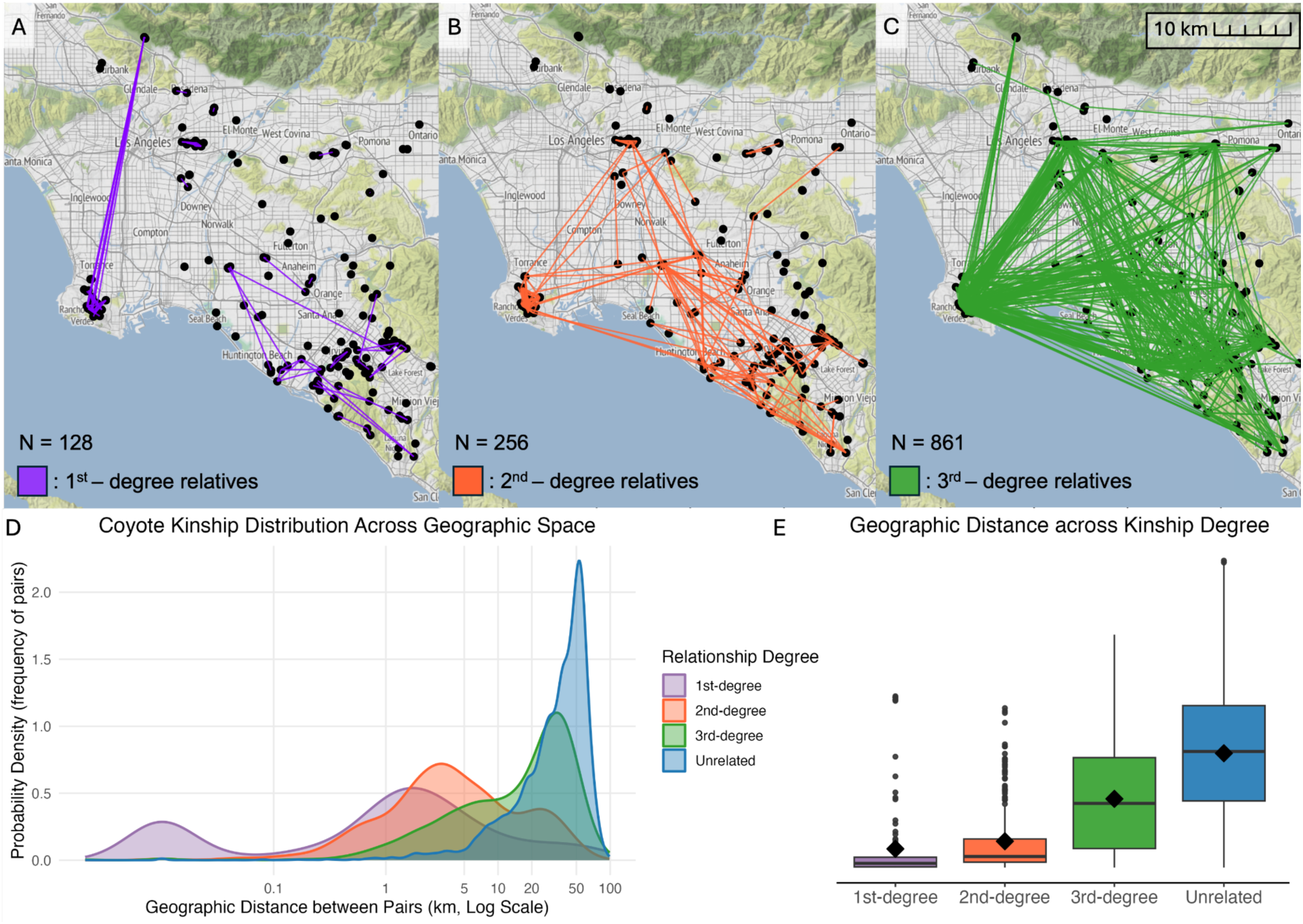
Spatial distribution of pairwise genomic kinship. **(A–C)** Geographic networks linking pairs of **(A)** first-degree (N = 128, orange), **(B)** second-degree (N = 256, purple), and **(C)** third-degree (N = 861, green) relatives across the study area. **(D)** Probability density distributions of pairwise geographic distances (km, log-scaled) partitioned by kinship category: first-degree (orange), second-degree (purple), third-degree (green), and unrelated pairs (blue). **(E)** Geographic distance distributions (km) across kinship tiers. Boxplots show the median (horizontal line), interquartile range (IQR; box limits), 1.5 x IQR (whiskers), mean values (black diamonds), and extreme outliers (individual points).

### Permeable highways facilitate asymmetric gene flow into the urban core

To explicitly assess dispersal across suspected anthropogenic barriers and interrogate the source-sink dynamics of specific geographic regions, we partitioned individuals manually into groups based on geographic location and ran two gene flow models: a “highway” model testing inferred barriers to migration (roadways I-5, I-10, I-405, and SR-60, where “I” stands for interstate, and “SR” stands for state route), and a “source-sink” model testing urban versus exurban groups.

The highway analysis consisted of six populations (Fig 3A): north of Interstate-10 (N I-10), in between I-10 and SR-60 (Inter I-10/SR-60), northeast of I-5 (NE I-5), in between I-5 and I-405 (Inter I-5/I-405), southwest of I-405 (SW 405), and the Palos Verdes Peninsula (PV; Figure 5d). The NE I-5 population served as a primary genetic source, exporting substantial directional gene flow into all other populations (Inter I-5/I-405 = 24.0%; Inter I-10/SR-60 = 22.8%; N I-10 = 22.9%, PV = 17.7%, and SW 405 = 13.7%). Additional notable asymmetric migration was observed from N I-10 into Inter I-10/SR-60 (5.9%) and from Inter I-5/I-405 into SW I-405 (8.77%), demonstrating permeable, non-random gene flow across major transportation corridors. Detailed migration rates for all pairwise comparisons are provided in Supplementary Table 25.

**Figure 3.**
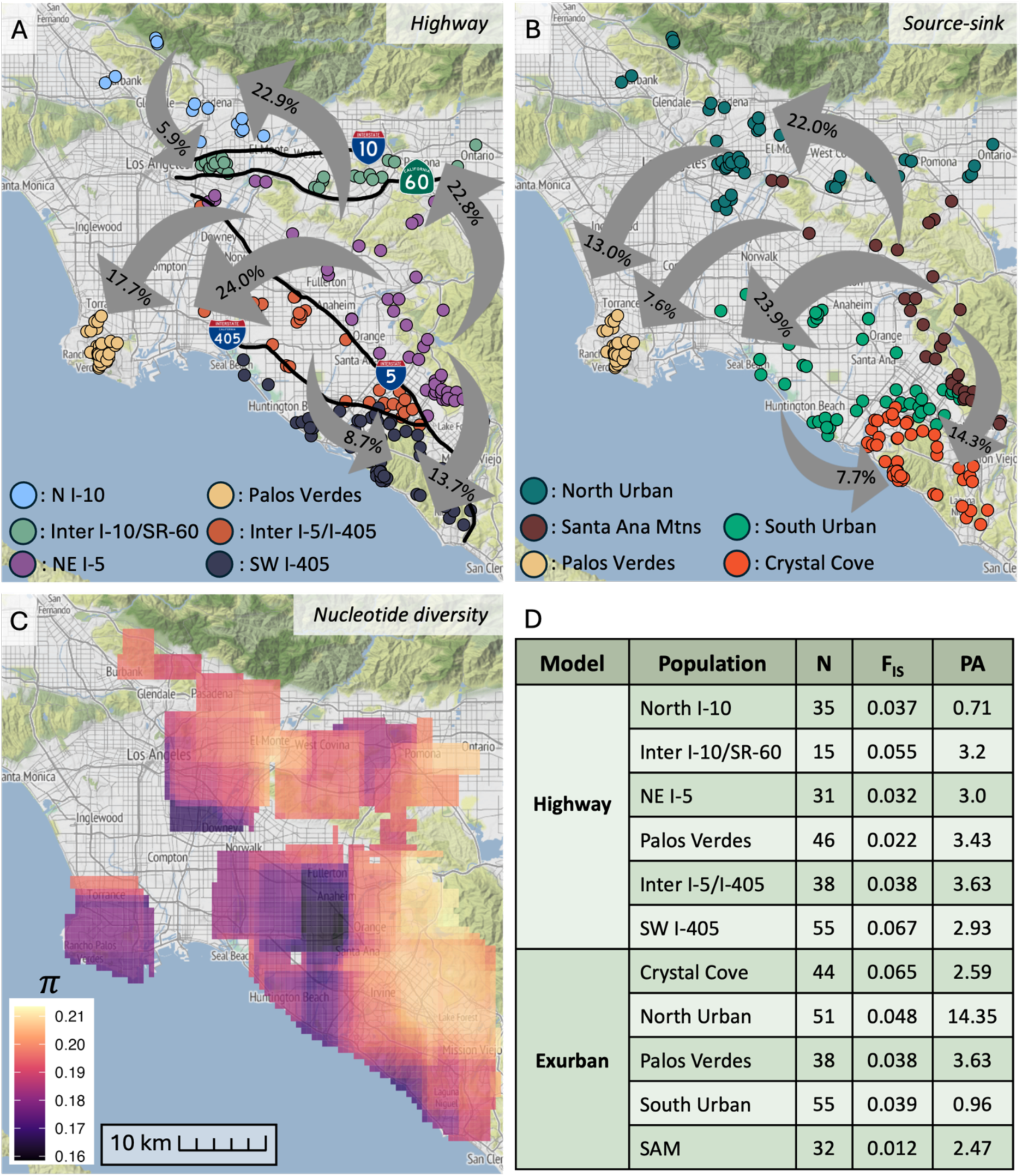
Contemporary gene flow models. **(A-B)** BayesAss3-SNPs inferred directional contemporary gene flow indicated via arrow, and quantified via reported percentage and arrow width. Circles denote individual sampling locations colored by manually assigned geographic population. Only rates >5% are visualized. **(A)** Highway Model: Inferred asymmetric migration rates (%) between six populations defined by major transportation corridors (I-5, I-10, I-405, and SR-60; represented by black lines). **(B)** Source-Sink Model: Migration rates (%) across five manually designated populations to test urban-exurban dynamics. **(C)** Spatial nucleotide diversity (pi) shown as a continuous spatial heatmap across the study area, with values ranging from lower diversity (dark purple) to higher diversity (light yellow). **(D)** Population Summary Table: Sample sizes (N), mean inbreeding coefficients (F_IS_), and normalized private allele counts (PA) calculated for all demographic units across both the Highways and Source-Sink spatial models.

We designated five populations in the source-sink model (Fig 3B): north urban, south urban, Santa Ana Mountains (SAM), Crystal Cove (CC), and the Palos Verdes Peninsula (PV). The two urban populations experienced significantly higher levels of impervious surface cover, pollution, and human and road density than the non-urban core populations (Figure S7). SAM was the chief source population, with substantial migration rates into all other populations (CC = 14.3%, PV = 7.6%, north urban = 22%, south urban = 23.9%) and very high philopatry (93.7%). The only other population to have an emigration rate >5% was south urban to CC (7.7%). PV, CC, and both urban sites had similar levels of philopatry (70.3-75.5%) and acted as net sinks in the landscape. Detailed migration rates for all pairwise comparisons are provided in Supplementary Table 26.

## Discussion

### Relatedness skews signatures of intra-urban contemporary evolution

A challenge in urban genomics is untangling true, population-level genetic structure from localized family-level dynamics. In broad-scale population genetics, sampling across large geographic scales minimizes the likelihood of sampling close kin (26). But within compressed urban landscapes, fine-scale sampling drastically increases the probability of capturing related individuals. This is especially true in pack animals like coyotes, which form tight-knit, territorial pack structures and social family units. In fragmented urban landscapes, these family groups often occupy localized habitat patches or resource hotspots, which can artificially inflate local genetic similarity (27,28). While sampling related individuals provides valuable insights into processes like contemporary dispersal and kin-biased movement, their inclusion in broader population genetic analyses can be problematic. Major genetic clustering algorithms are highly sensitive to family structure and rely on the assumption of unrelated samples (29–31). Thus, failing to account for close kin inherently biases population assignment. Prior to kinship filtering, our unfiltered dataset (N = 220) suggested five distinct genetic clusters. However, removing first-degree relatives collapsed this structure to three geographically coherent populations. Removing up to second-degree relatives rendered one to three clusters equally likely, underscoring that once all family structure is stripped away, population structure is quite weak.

Kin structure is a known driver of fine-scale genetic patterns in social mammals (28,32), but handling relatedness remains a point of significant inconsistency across the urban wildlife literature. Some coyote studies filter first-degree relatives as a standard precaution (8), report that relatedness does not meaningfully alter clustering (33), intentionally retain kin to analyze social pack dynamics (28), or do not report relatedness at all (34,35). We suspect the conclusion that relatedness does not affect population assignment is largely a methodological artifact of genetic marker; earlier studies relied on microsatellite panels, which often lack the statistical power to resolve fine-scale family clusters particularly amidst high background gene flow (36). By leveraging RADSeq to generate thousands of genome-wide SNPs, our dataset possessed the resolution required to detect tight familial ties and their influence on population structure, which may not have been detectable with microsatellites. While retaining relatives is appropriate when explicitly investigating fine-scale social dynamics or pack pedigree, doing so in macro-population frameworks risks conflating localized family packs with isolated, genetically divergent subpopulations. Consequently, these results highlight that unmitigated kinship structure can distort population inference, underscoring the need for explicit kinship filtering when assessing regional connectivity in social mammals.

Across the three coyote populations that did hold with first and second-degree relatives removed, we observed low-to-moderate genetic differentiation characteristic of highly mobile urban carnivores capable of long-distance dispersal (33,37–40). However, the presence of private alleles across all populations indicates sufficient isolation for unique genetic signatures to persist (41,42). We detected signatures of mild inbreeding overall, though more intense inbreeding occurred in a small subset of individuals. However, levels of genetic diversity remained robust, suggesting that urban fragmentation, though driving some genetic structuring, has not reached a threshold of critical genetic erosion.

Notably, these population genetic results alone could represent a snapshot of either diverging or converging subpopulations. Given that neutral evolutionary processes typically operate over thousands of years, static spatial sampling cannot easily distinguish between ongoing divergence driven by urban barriers and convergence driven by corridor use in a very young metropolis like Greater LA. Resolving this uncertainty would require integrating longitudinal monitoring and/or historical samples to explicitly model temporal genetic change across the urban matrix. However, because the Los Angeles Basin historically provided continuous habitat occupied by coyotes since the Late Pleistocene (37,38), we can assume natural landscape barriers were minimal. The primary hydrological feature, the Los Angeles River, was certainly navigable by coyotes, though it could have delineated territorial or social pack boundaries. Coyote populations pre-settlement and development across this study area were therefore likely broadly panmictic, especially given our sampling excludes the region’s major topographically isolating Santa Monica and San Gabriel mountain ranges. Thus, we can infer that the observed genetic structuring, while weak, is likely driven by modern urban infrastructure on an accelerated timescale due to urban selective pressures and may become more pronounced over time (1).

### Mapping relatedness acts as a proxy for dispersal and reveals a permeable landscape

Spatial mapping of close kin provides a powerful genomic proxy for direct dispersal events, capturing realized long-distance gene flow without the logistical constraints of animal tracking. This approach may be particularly valuable in urban areas, where animal movement studies remain rare (45). In particular, mapping first degree relatives acts as a proxy for real dispersal events of parent-offspring or sibling-sibling dispersal. We identified several first-degree relative pairs >50 km apart, representing remarkably long-range dispersal events. While these long-distance linkages primarily reflect individual dispersal, there is preliminary evidence that suggests whole family groups may occasionally relocate or disperse together across disturbed landscapes - a phenomenon that could also yield distant clusters of close kin (27). Regardless of the underlying movement mechanism, this finding is particularly striking given urban coyote spatial ecology work. In urban environments, mammalian movement is generally heavily constrained with global trends suggesting that movement extents in high-human-footprint areas are reduced to one-half or one-third of those in rural landscapes (46). Telemetry work in Toronto confirmed urban coyotes showed significantly less long-range movement than their rural counterparts (47). Researchers similarly found urban coyotes in Chicago had attenuated dispersal distances compared to ranges reported in rural coyotes (48). However, they also demonstrated that coyotes originating from more developed natal habitats were more likely to disperse, and covered greater distances, than those raised in less developed areas (48). Integrating spatial genomics with empirical movement data is therefore essential for fully resolving urban dispersal dynamics, as genetic linkages capture realized gene flow events that tracking alone may miss.

### Landscape genetics models support the urban facilitation model: coyotes experience few anthropogenic genetic barriers

Contrary to our expectations, we found no significant relationship between six environmental variables and any metric of diversity (Ho, Fis, allelic richness, private alleles). Genetic diversity has been found to linearly decrease with increasing urbanization, specifically in coyotes (37,38) and more broadly across mammals (7). Here, we attempted to tease apart what within the urban environment may be driving this pattern. We characterized fine-scale environmental variables including human population density, impervious surface cover, NDVI, pollution burden, road density, and a time-series dataset of historical built-up intensity. By including human population density and pollution burden we highlight social-ecological relationships, which are often excluded from evolutionary work (19). The ability to classify different ecosystems at the intra-urban scale highlights the importance of moving away from an urban-rural dichotomy and single “all-encompassing” urbanization metrics (49). Ultimately, the lack of correlation between these environmental trends and genetic statistics underscores the necessity of transitioning from simple linear comparisons to a more nuanced landscape genomics framework. Such models are better equipped to capture the complex, non-linear mechanisms through which the urban matrix structures contemporary gene flow (50).

Even when utilizing high-resolution landscape genetics models, we were unable to identify specific urban environmental drivers of genetic differentiation. Despite rigorously testing IBE and IBR models with our urban variables of interest, only geographic distance (IBD) significantly correlated with genetic differentiation. A pattern of IBD is not surprising, given animals have a limited dispersal capacity, which may be further restricted across highly populated and urbanized landscapes (51). However, we anticipated that natal-biased habitat dispersal may contribute to a pattern of IBE, which has been identified in coyotes across Central California ecoregions (52). Additionally, some urban coyotes within Chicago show patterns of natal-habitat biased dispersal (48). Environmental factors not only contribute to stressor and pollutant exposure, but also to prey availability, food subsidies, mating opportunities, etc. all of which can contribute to environmental specialization (53) and thus a pattern of IBE. However, given the relatively fine-scale variation between our urban niches along with our migration results suggesting maintained connectivity across regions, the lack of a significant IBE signal is perhaps unsurprising.

We expected IBR would outperform models of IBE, as resistance models explicitly account for the physical landscape features that individuals must navigate during dispersal to achieve functional genetic connectivity. In addition, IBR has more often been the focus of landscape genetics studies in anthropogenically altered environments (39,54–58). Studies have identified tree cover (54), impervious surface cover (39), roads/linear barriers (55), and general urbanization metrics (56) to be driving genetic processes in various mammals. However, our analyses did not reach significance across any metric: roads as barriers, roads as corridors, and NDVI each run at three resistance magnitudes (low, medium, and high). While a single negative finding may be expected, the total absence of significant IBR or IBE models across all tested variables is striking. This lack of environmental or resistance-driven structure underscores the coyote’s remarkable behavioral plasticity and high dispersal capability even within an extremely dense urban matrix.

We further utilized HISDAC-US time-series data (59) to test whether contemporary genetic structure was a product of historical landscape configurations using both IBE and IBR models. We hypothesized that the rapid mid-century expansion of the urban matrix and particularly the freeway system would produce a discrete peak at that time in the explanatory power of our models. Alternatively, a saturation curve in explanatory power could indicate a baseline level of urbanization, rather than specific development, dictated current structure. However, no historical model reached significance for any decade. This lack of temporal signal suggests that contemporary coyote genetic patterns are not anchored to historical development phases in this landscape. Instead, the high vagility and behavioral plasticity of the species likely allow for ongoing, high-frequency movement that overrides the genetic signatures of past landscapes. As mentioned previously, resampling these populations over the coming decades will reveal whether fine-scale genetic divergence accelerates over time or remains buffered by continuous, plastic movement through the urban matrix. This is particularly relevant given the very recent development of Greater LA on an evolutionary timescale.

Our results present a complex evolutionary paradox: while the lack of significant IBE and IBR suggests the urban matrix is permeable to coyote movement, the presence of population structure indicates a fragmented genetic landscape. This apparent contradiction suggests that non-structural, behavioral factors may be limiting effective dispersal. Riley et al. 2006 observed coyote territorial barriers along highways, maintaining genetic gaps even when dispersal across the highway was successful. Given our lack of IBR finding here, these territorial barriers likely exist even in the absence of physical obstructions. We propose that coyote density may be a critical, untested driver of this pattern, where high-density, saturated habitats impose “social resistance” that forces dispersers to move through territories without successfully establishing residency. Furthermore, if high urban coyote densities allow individuals to find mates in close proximity, the resulting shortened dispersal distances would naturally drive the observed signature of IBD.

While our IBE and IBR models did not result in significant findings, we strongly recommend continuing to apply these rigorous models to urban evolution work. Incorporating fine-scale social-ecological and infrastructure data, high-resolution sequencing data, and historical landscape data into landscape genomics models provides a scalable approach for studying the nuanced drivers of urban evolution. Applying these methods to less vagile species, species with shorter generation times, and/or populations inhabiting older urban centers may reveal stronger, more readily detectable drivers of urban-driven evolution.

### Gene flow analyses further support urban permeability finding for coyotes

Substantial migration rates were detected across I-5, I-405, I-10, and SR-60. These results indicate that despite the high cost of crossing these roadways, successful dispersers are frequent enough to facilitate robust gene flow, which further supports our landscape genetics models’ null findings. This aligns with findings in a Central Mexican coyote population (57) but contradicts a previous study in Southern California that identified highways as drivers of genetic fragmentation (60). However, neither of these studies explicitly calculated gene flow rates nor used next-generation sequencing, which could explain the inconsistency. Urban genetic work explicitly measuring contemporary gene flow is rare and focused on rural-urban (61,62), inter-city (54,63), or mixed landuse, wider geographic (56,64) dispersal. Quantifying intra-urban migration rates between distinct metropolitan sub-populations is important as it shifts the focus from inferring movement through static resistance models to directly measuring functional connectivity.

Crucially, these results suggest a decoupling of ecological and genetic barriers; while landscape features such as high-traffic roadways undoubtedly function as ecological barriers that impose significant individual mortality, they do not function as genetic barriers. Furthermore, our sample location represents each coyote’s place of death. We use this point to estimate home range, but individuals may have been actively dispersing when killed. We attempted to conduct additional gene flow assessments removing roadkill individuals to correct for an overrepresentation of dispersing individuals, but did not have enough sampling density to perform robust analysis after they had been removed.

### Source-sink dynamics support a “leaky sink” model

Our results do not support a strict urban sink model, wherein urban patches act purely as terminal genetic repositories, but instead demonstrate that urban core coyote populations in this landscape also export gene flow. The two urban core populations receive substantial directional gene flow from the regional source population (Santa Ana Mountains) with nearly no back-migration. However, these urban groups then actively pass genetic material further westward into less urban habitat (Palos Verdes Peninsula and Crystal Cove) while maintaining stable levels of genetic diversity. Further, the northern urban core harbors an unusually high abundance of private alleles, operating as a “genetic sponge” that absorbs diverse regional variation via immigration. Notably, the urban cores have near-zero immigration rates with each other, perhaps suggesting saturated habitat in the highest density urban areas. Notably, immigration between the two urban cores is virtually absent, suggesting that localized habitat saturation may restrict direct movement between adjacent urban centers.

We expected the less developed (higher NDVI, lower population density) habitats on the coast to act as regional sources. Rather, they both functioned as sinks with minimal emigration. The Crystal Cove population, though encompassing a large, undeveloped state park and highly dispersed housing, had the highest inbreeding levels and lowest diversity. Importantly, these demographic disruptions and directional dispersal patterns are likely intensified, if not driven, by lethal control management. Wildlife populations in urban cores and affluent suburban areas experience lethal management (65). Frequent coyote removal can create territory vacancies, which draw dispersers. Removal events also repeatedly fracture multi-generational pack structures, and surviving local individuals may be forced to mate with close relatives between turnover events, driving up inbreeding while simultaneously maintaining a high net influx of wildland migrants. Finally, dispersal out of both Crystal Cove and the Palos Verdes Peninsula is geographically restricted to the west by the Pacific Ocean, transforming these coastal fragments into the terminal demographic traps of this study area.

Source–sink dynamics remain poorly resolved across urban ecosystems (66). Historically, studying source-sink dynamics was generally difficult due to the extensive monitoring needed to follow individuals and quantify their reproductive success (67,68). However, modern high-throughput genomic tools provide a scalable framework to infer these fine-scale movement dynamics. In mammals, urban source–sink patterns vary widely depending on species life history and localized matrix features. For instance, Virginia opossums (*Didelphis virginiana*) leveraging urban resource subsidies acted as demographic sources for adjacent natural areas (69). Meanwhile, wild boars (*Sus scrofa*) in Berlin exhibited classic urban sink dynamics (70). Black bears (*Ursus americanus*) and pumas (*Puma concolor*) similarly exhibited sink populations in more developed landscapes, though these were largely situated along exurban fringes (71–73). Because these earlier studies relied primarily on lower-density markers like microsatellites, transitioning to genome-wide SNP datasets is critical for resolving fine-scale, contemporary movement with greater statistical power. Crucially, source–sink relationships are not static; they represent fluid demographic processes that can shift dynamically across both seasonal and long-term evolutionary timescales (66,68,69).

## Conclusion

Our findings demonstrate that integrating fine-scale genomic resolution with landscape modeling that incorporates social-ecological variables can reshape paradigms in urban evolutionary biology. By illustrating how unmitigated kin structure can artificially mimic genetic differentiation, we establish the critical need to disentangle local pedigree dynamics from macro-population structure when evaluating anthropogenic fragmentation. We demonstrate that highly plastic generalists can maintain functional genetic connectivity across hyper-fragmented megacities, and that population structure is driven primarily by social resistance, isolation-by-distance, and anthropogenic mortality rather than landscape features or habitat characteristics. Finally, our results demonstrate a striking nuance to the traditional urban sink model by revealing a decoupling between demographic and genetic sink dynamics. Demographically, the urban core functions as a classic sink: it receives heavy directional immigration from the wildland source with negligible back-migration, while high urban mortality and fragmentation drive elevated local inbreeding. Genetically, however, the urban core does not act as a terminal dead end. Instead, it functions as a “leaky sink,” exporting sufficient genetic diversity westward to prevent inbreeding depression in coastal terminal patches. Furthermore, one urban core population functions as a “genetic sponge,” actively absorbing and harboring transient alleles from multiple surrounding source populations. Thus, urban habitat can simultaneously exhibit sink-like demographic traits while maintaining functional genetic connectivity and even genetic novelty across heavily altered landscapes. As urbanization accelerates globally, applying this multi-tiered genomic framework will be essential for predicting how wildlife populations navigate, adapt to, and persist within the world’s rapidly expanding developed landscapes.

## Materials & Methods

### Sample Collection and Genomic Sequencing

We opportunistically collected tissue samples from coyote roadkill carcasses and agency management removals between 2016 and 2022 across Los Angeles, Orange, San Bernardino, Riverside, and San Diego counties (*SI Appendix*). Genomic DNA was extracted using the Qiagen DNeasy Blood and Tissue Kit and quantified via Qubit fluorometry. Double-digest RADseq (ddRADseq) libraries were constructed using the bestRAD protocol with the *SfbI*-HF restriction enzyme. Adapter-ligated fragments were enriched, pool-sonicated to 300–400 bp on a Covaris LE220, and indexed with dual-barcode Illumina P2 adapters. Paired-end sequencing (2 x 150 bp) was performed on an Illumina NovaSeq 6000 platform at the Yale Center for Genome Analysis, yielding approximately 4 million reads per sample.

### Bioinformatic Processing and Variant Filtering

Raw reads were processed using STACKS v2.6 (74). Barcodes and low-quality reads (quality score <10) were filtered and demultiplexed via *process_radtags* with up to 2bp barcode mismatches permitted. PCR duplicates were removed using *clone_filter.* Retained reads were aligned to the dog reference genome (76; CanFam3.1) supplemented with the domestic dog Y chromosome (76) using bwa-mem v0.7.19 (77). Variant calling was executed using the Maruki-Low model in STACKS (gstacks and populations, --vt-alpha and --gt-alpha with p = 0.01). High-confidence SNPs were retained using VCFtools v0.1.17 (78) by removing loci with >10 missing data, singletons, and private doubletons, while filtering out individuals with >20 missing genotypes. PLINK2 (79) was used to apply minor allele frequency filtering (MAF <0.03). For demographic analyses that require neutral and unlinked loci, we constructed a “demographic dataset” by excluding loci in linkage disequilibrium (LD) using a moderate genotype correlation threshold (r^2^>0.5) in *PLINK2* (--indep-pairwise 50 5 0.2) and loci that significantly deviated from Hardy-Weinberg Equilibrium (--hwe, *p*=0.001). For calculating relatedness, we applied additional filtering to retain loci with MAF>0.4 and lower levels of missing data, permitting only 5% missing data (*SI Appendix*).

### Population Genetics, Kinship, and Spatial Diversity

Population structure was evaluated using Principal Component Analysis (PCA) and ADMIXTURE v1.3.0 (29; K = 1–10, 10 cross-validation replicates per K). Consensus Q-matrices were computed using CLUMPP v1.1.2 (80). To disentangle macro-population structure from local family dynamics, pairwise kinship coefficients were calculated using the KING algorithm in PLINK2 (79). Analyses were repeated after removing first-degree (k > 0.177) and second-degree (k > 0.0884) relatives. Population differentiation was quantified using Weir and Cockerham’s F_ST i_n VCFtools (78). Individual observed (H_O)_ and expected (H_E)_ heterozygosity and inbreeding coefficients (F_IS)_ were estimated in VCFtools, while standardized allelic richness (AR) and private allele counts (PA) were calculated in R using *hierfstat* and *poppr (81,82)*.

Continuous spatial surfaces of nucleotide diversity (pi) were generated using moving-window ordinary kriging in *wingen* v1.0 (83).

### Quantification of Social-Ecological Covariates

To evaluate environmental drivers of gene flow, we quantified five social-ecological parameters: human population density, fractional impervious surface cover, Normalized Difference Vegetation Index (NDVI), pollution burden, and road density. To account for spatial sampling uncertainty around mortality locations, each GPS point was buffered by a 1km radius for a home range proxy (3.14km^2^; Poisson et al.). Human population density and road density were derived at the census tract level using 2020 U.S. Census data and the R package *tigris* v2.1 (85). Impervious surface cover was extracted from the 2020 NLCD dataset (30 m resolution). Springtime NDVI was derived from Landsat 8 surface reflectance data (<10% cloud cover). A cumulative pollution burden index was calculated from CalEnviroScreen 4.0 (86) combining five exposure parameters (lead, particulate matter, diesel particulate matter, toxic releases, solid waste) and four environmental effect parameters (groundwater threat, hazardous waste, cleanup sites, impaired water bodies). All covariates were normalized (0–100), and pairwise collinearity was assessed using Pearson correlation tests (r < 0.70).

### Landscape Genetics and Resistance Modeling

Associations between genetic diversity metrics (H_O,_ F_IS,_ allelic richness, private alleles) and urban covariates were evaluated using Spearman’s rank correlations (rho). Isolation by Distance (IBD) was tested using Multiple Matrix Regression with Randomizations (MMRR, 1,000 permutations). Isolation by Environment (IBE) was evaluated using Generalized Dissimilarity Modeling (GDM) implemented in the R package *algatr* v1.0 (87). Isolation by Resistance (IBR) was modeled using Circuitscape v5.15.0 in pairwise mode (88). Resistance surfaces were parameterized for NDVI and road density both with roads as corridors and barriers (all at 150m resolution). Resistance values were tested under low, moderate, and high transformation functions. Partial Mantel tests evaluated pairwise resistance matrices against genetic dissimilarity while controlling for Euclidean distance. To test for lag effects from historical urbanization, time-series Built-Up Intensity (BUI) layers from HISDAC-US (59; 1940–2000, 5-year increments, 250m resolution) were modeled in GDM and Circuitscape. Fine-scale spatial pedigrees were mapped using first- and second-degree relative pairs to track contemporary trans-barrier dispersal events.

### Contemporary Directional Gene Flow

Directional migration rates over recent generations were estimated using BayesAss3 (BA3-SNPs v3.0.4; 73) on the unlinked dataset of 1,464 loci. We parameterized two distinct spatial frameworks: a "source-sink" model testing migration between urban and exurban cohorts, and a "highway" model evaluating connectivity across major transportation corridors (I-5, I-10, I-405, and SR-60). MCMC mixing parameters were optimized using BA3-SNPs-autotune v2.1.2 (89). Five independent chains per model were executed with different random seeds for 20,000,000 iterations (5,000,000 burn-in) for the source-sink model. The highway model took longer to converge with 50,000,000 iterations (10,000,000 burn-in). Chain convergence was confirmed via MCMC trace plots in R (*SI Appendix*, Fig. S8), and posterior migration rates were calculated as the mean standard deviation across independent replicates.

## Citations

1. Johnson MTJ, Munshi-South J. Evolution of life in urban environments. Science. 2017 Nov 3;358(6363):eaam8327. doi:10.1126/science.aam8327

2. Simkin RD, Seto KC, McDonald RI, Jetz W. Biodiversity impacts and conservation implications of urban land expansion projected to 2050. Proc Natl Acad Sci. 2022 Mar 22;119(12):e2117297119. doi:10.1073/pnas.2117297119

3. Griffith J, Sunday JM, Hargreaves AL. Urbanization weakens the global latitudinal diversity gradient in birds. Proc Natl Acad Sci U S A. 2026 Aug 25;123(34):e2527860123. doi:10.1073/pnas.2527860123 PubMed PMID: 42612001.

4. Miles LS, Rivkin LR, Johnson MTJ, Munshi-South J, Verrelli BC. Gene flow and genetic drift in urban environments. Mol Ecol. 2019 Sep;28(18):4138–51. doi:10.1111/mec.15221

5. Verrelli BC, Alberti M, Des Roches S, Harris NC, Hendry AP, Johnson MTJ, et al. A global horizon scan for urban evolutionary ecology. Trends Ecol Evol. 2022 Nov;37(11):1006–19. doi:10.1016/j.tree.2022.07.012

6. McCluney KE, Deviche P, Sweazea KL, Carlen EJ, Clark JAG, Grade AM, et al. An integrated social–ecological–evolutionary–phenotypic (SEEP) approach to understanding animal responses to urbanization. Biol Rev. 2026 Feb;101(1):419–36. doi:10.1111/brv.70088

7. Schmidt C, Domaratzki M, Kinnunen RP, Bowman J, Garroway CJ. Continent-wide effects of urbanization on bird and mammal genetic diversity. Proc R Soc B Biol Sci. 2020 Feb 12;287(1920):20192497. doi:10.1098/rspb.2019.2497

8. Adducci A, Jasperse J, Riley S, Brown J, Honeycutt R, Monzón J. Urban coyotes are genetically distinct from coyotes in natural habitats. J Urban Ecol. 2020 Jan 1;6(1):juaa010. doi:10.1093/jue/juaa010

9. Serieys LEK, Jackson M, Sleater-Squires S, Leighton GRM, Drouilly M, Viljoen S, et al. Urbanization drives genetic erosion and population structure in a historically connected carnivore population. Conserv Genet. 2026 Apr;27(2):46. doi:10.1007/s10592-026-01776-9

10. Pulliam HR. Sources, sinks, and population regulation. Am Nat. 1988 Nov;132(5):652–61.

11. Battin J. When Good Animals Love Bad Habitats: Ecological Traps and the Conservation of Animal Populations. Conserv Biol. 2004 Dec;18(6):1482–91. doi:10.1111/j.1523-1739.2004.00417.x

12. Lepczyk CA, Aronson MFJ, Evans KL, Goddard MA, Lerman SB, MacIvor JS. Biodiversity in the City: Fundamental Questions for Understanding the Ecology of Urban Green Spaces for Biodiversity Conservation. BioScience. 2017 Sep;67(9):799–807. doi:10.1093/biosci/bix079

13. Luck GW. A review of the relationships between human population density and biodiversity. Biol Rev. 2007 Nov;82(4):607–45. doi:10.1111/j.1469-185X.2007.00028.x

14. Mcdonald RI, Kareiva P, Forman RTT. The implications of current and future urbanization for global protected areas and biodiversity conservation. Biol Conserv. 2008 Jun;141(6):1695–703. doi:10.1016/j.biocon.2008.04.025

15. Wilkinson CE, Quinn N, Eng C, Schell CJ. Environmental Health and Societal Wealth Predict Movement Patterns of an Urban Carnivore. Ecol Lett. 2025 Feb;28(2):e70088. doi:10.1111/ele.70088

16. Hill J, DeVault T, Belant J. Comparative influence of anthropogenic landscape pressures on cause-specific mortality of mammals. Perspect Ecol Conserv. 2022 Jan;20(1):38–44. doi:10.1016/j.pecon.2021.10.004

17. Hill JE, DeVault TL, Belant JL. Cause-specific mortality of the world’s terrestrial vertebrates. Algar A, editor. Glob Ecol Biogeogr. 2019 May;28(5):680–9. doi:10.1111/geb.12881

18. Pickett STA, Burch Jr. WR, Dalton SE, Foresman TW, Grove JM, Rowntree R. A conceptual framework for the study of human ecosystems in urban areas. Urban Ecosyst. 1997;1(4):185–99. doi:10.1023/A:1018531712889

19. Des Roches S, Brans KI, Lambert MR, Rivkin LR, Savage AM, Schell CJ, et al. Socio-eco-evolutionary dynamics in cities. Evol Appl. 2021 Jan;14(1):248–67. doi:10.1111/eva.13065

20. Estien C, Fidino M, Wilkinson C, Morello-Frosch R, Schell C. Historical redlining impacts wildlife biodiversity across California [Internet]. Ecology and Evolutionary Biology; 2023 [cited 2025 Dec 19]. Available from: https://ecoevorxiv.org/repository/view/6378/doi:10.32942/X24K60

21. Hentati Y, Estien CO, Hawn Z, Jordan MJ, Long RA, Mueller R, et al. Environmental contamination predicts mammal diversity and mesocarnivore activity in the Seattle-Tacoma metro area. Urban Ecosyst. 2025 Aug;28(4):152. doi:10.1007/s11252-025-01758-8

22. Ellis-Soto D, Chapman M, Locke DH. Historical redlining is associated with increasing geographical disparities in bird biodiversity sampling in the United States. Nat Hum Behav. 2023 Sep 7;7(11):1869–77. doi:10.1038/s41562-023-01688-5

23. Schell CJ, Dyson K, Fuentes TL, Des Roches S, Harris NC, Miller DS, et al. The ecological and evolutionary consequences of systemic racism in urban environments. Science. 2020 Sep 18;369(6510):eaay4497. doi:10.1126/science.aay4497

24. Moran PA, Bosse M, Mariën J, Halfwerk W. Genomic footprints of (pre) colonialism: Population declines in urban and forest túngara frogs coincident with historical human activity. Mol Ecol. 2024 Feb;33(4):e17258. doi:10.1111/mec.17258

25. Epps CW, Keyghobadi N. Landscape genetics in a changing world: disentangling historical and contemporary influences and inferring change. Mol Ecol. 2015 Dec;24(24):6021–40. doi:10.1111/mec.13454

26. Östergren J, Palm S, Gilbey J, Dannewitz J. Close relatives in population samples: Evaluation of the consequences for genetic stock identification. Mol Ecol Resour. 2020 Mar;20(2):498–510. doi:10.1111/1755-0998.13131 PubMed PMID: 31883430; PubMed Central PMCID: PMC7065253.

27. Bird S, Monzón JD, Meyer WM, Moore JE. An Illusion of Barriers to Gene Flow in Suburban Coyotes (Canis latrans): Spatial and Temporal Population Structure across a Fragmented Landscape in Southern California. Diversity. 2023 Apr 1;15(4):498. doi:10.3390/d15040498

28. Henger CS, Herrera GA, Nagy CM, Weckel ME, Gormezano LJ, Wultsch C, et al. Genetic diversity and relatedness of a recently established population of eastern coyotes (Canis latrans) in New York City. Urban Ecosyst. 2020 Apr;23(2):319–30. doi:10.1007/s11252-019-00918-x

29. Alexander DH, Novembre J, Lange K. Fast model-based estimation of ancestry in unrelated individuals. Genome Res. 2009 Sep;19(9):1655–64. doi:10.1101/gr.094052.109

30. Hubisz MJ, Falush D, Stephens M, Pritchard JK. Inferring weak population structure with the assistance of sample group information. Mol Ecol Resour. 2009 Sep;9(5):1322–32. doi:10.1111/j.1755-0998.2009.02591.x

31. Lawson DJ, Van Dorp L, Falush D. A tutorial on how not to over-interpret STRUCTURE and ADMIXTURE bar plots. Nat Commun. 2018 Aug 14;9(1):3258. doi:10.1038/s41467-018-05257-7

32. Parreira BR, Chikhi L. On some genetic consequences of social structure, mating systems, dispersal, and sampling. Proc Natl Acad Sci. 2015 Jun 30;112(26). doi:10.1073/pnas.1414463112

33. Kreling SES, Reese EM, Cavalluzzi OM, Bozzi NB, Messinger R, Schell CJ, et al. City divided: Unveiling family ties and genetic structuring of coyotes in Seattle. Mol Ecol. 2024 Jul;33(14). doi:10.1111/mec.17427

34. DeCandia AL, Henger CS, Krause A, Gormezano LJ, Weckel M, Nagy C, et al. Genetics of urban colonization: neutral and adaptive variation in coyotes (*Canis latrans*) inhabiting the New York metropolitan area. J Urban Ecol. 2019 Jan 1;5(1). doi:10.1093/jue/juz002

35. Rebecca M. Rashleigh, Robert A. Krebs, Harry Van Keulen. Population structure of coyote (Canis latrans) in urban landscape (Cleveland, Ohio, USA). Ohio J Sci. 2008 Sep;108(4).

36. Lemopoulos A, Prokkola JM, Uusi-Heikkilä S, Vasemägi A, Huusko A, Hyvärinen P, et al. Comparing RADseq and microsatellites for estimating genetic diversity and relatedness -Implications for brown trout conservation. Ecol Evol. 2019 Feb;9(4):2106–20. doi:10.1002/ece3.4905 PubMed PMID: 30847096; PubMed Central PMCID: PMC6392366.

37. Adducci A, Jasperse J, Riley S, Brown J, Honeycutt R, Monzón J. Urban coyotes are genetically distinct from coyotes in natural habitats. J Urban Ecol. 2020 Jan 1;6(1):juaa010. doi:10.1093/jue/juaa010

38. DeCandia AL, Henger CS, Krause A, Gormezano LJ, Weckel M, Nagy C, et al. Genetics of urban colonization: neutral and adaptive variation in coyotes (*Canis latrans*) inhabiting the New York metropolitan area. J Urban Ecol. 2019 Jan 1;5(1). doi:10.1093/jue/juz002

39. Kozakiewicz CP, Burridge CP, Funk WC, Salerno PE, Trumbo DR, Gagne RB, et al. Urbanization reduces genetic connectivity in bobcats (*Lynx rufus*) at both intra–and interpopulation spatial scales. Mol Ecol. 2019 Dec;28(23):5068–85. doi:10.1111/mec.15274

40. Marshall CA, Halo J, Van Why K, Loera Y, Tennenbaum SR, Burton C, et al. Signatures of the Anthropocene: Population Genomic Structure Detected in Pennsylvania Coyotes. Ecol Evol. 2026 Mar;16(3):e73216. doi:10.1002/ece3.73216

41. Barton NH, Slatkin M. A Quasi-equilibrium theory of the distribution of rare alleles in a subdivided population. Heredity. 1986 Jun;56(3):409–15. doi:10.1038/hdy.1986.63

42. Slatkin M. RARE ALLELES AS INDICATORS OF GENE FLOW. Evolution. 1985 Jan;39(1):53–65. doi:10.1111/j.1558-5646.1985.tb04079.x

43. Dan Flores. Coyote America: A Natural and Supernatural History. Basic Books; 2016.

44. DeSantis LRG, Crites JM, Feranec RS, Fox-Dobbs K, Farrell AB, Harris JM, et al. Causes and Consequences of Pleistocene Megafaunal Extinctions as Revealed from Rancho La Brea Mammals. Curr Biol. 2019 Aug;29(15):2488–2495.e2. doi:10.1016/j.cub.2019.06.059

45. Ellis-Soto D, Flack A, Strandburg-Peshkin A, Wild TA, Williams HJ, O’Mara MT. From biologging to conservation: Tracking individual performance in changing environments. Proc Natl Acad Sci U S A. 2025 Aug 5;122(31):e2410947122. doi:10.1073/pnas.2410947122 PubMed PMID: 40720657; PubMed Central PMCID: PMC12337290.

46. Tucker MA, Böhning-Gaese K, Fagan WF, Fryxell JM, Van Moorter B, Alberts SC, et al. Moving in the Anthropocene: Global reductions in terrestrial mammalian movements. Science. 2018 Jan 26;359(6374):466–9. doi:10.1126/science.aam9712

47. Gelmi-Candusso TA, Wheeldon TJ, Patterson BR, Fortin MJ. The effect of urbanization and behavioral factors on coyote net displacement and its implications for seed dispersal. Urban Ecosyst. 2024 Apr;27(2):387–97. doi:10.1007/s11252-023-01460-7

48. Zepeda E, Payne E, Wurth A, Sih A, Gehrt S. Early life experience influences dispersal in coyotes (*Canis latrans*). Candolin U, editor. Behav Ecol. 2021 Aug 19;32(4):728–37. doi:10.1093/beheco/arab027

49. Pickett STA, Zhou W, Childers DL, Grove JM, Hansen WD, Locke DH, et al. The continuum of urbanity: a synthetic concept for research on urban-rural mixtures. Npj Urban Sustain. 2026 Jan 29;6(1):36. doi:10.1038/s42949-026-00347-8

50. Fusco NA, Carlen EJ, Munshi-South J. Urban Landscape Genetics: Are Biologists Keeping Up with the Pace of Urbanization? Curr Landsc Ecol Rep. 2021 Jun;6(2):35–45. doi:10.1007/s40823-021-00062-3

51. Meirmans PG. The trouble with isolation by distance. Mol Ecol. 2012 Jun;21(12):2839–46. doi:10.1111/j.1365-294X.2012.05578.x

52. Sacks BN, Brown SK, Ernest HB. Population structure of California coyotes corresponds to habitat-specific breaks and illuminates species history. Mol Ecol. 2004 May;13(5):1265–75. doi:10.1111/j.1365-294X.2004.02110.x

53. Caspi T, Johnson JR, Lambert MR, Schell CJ, Sih A. Behavioral plasticity can facilitate evolution in urban environments. Trends Ecol Evol. 2022 Dec;37(12):1092–103. doi:10.1016/j.tree.2022.08.002

54. Fusco NA, Cosentino BJ, Gibbs JP, Allen ML, Blumenfeld AJ, Boettner GH, et al. Population genomic structure of a widespread, urban-dwelling mammal: The eastern grey squirrel (*Sciurus carolinensis*). Mol Ecol. 2024 Feb;33(3):e17230. doi:10.1111/mec.17230

55. Kimmig SE, Beninde J, Brandt M, Schleimer A, Kramer-Schadt S, Hofer H, et al. Beyond the landscape: Resistance modelling infers physical and behavioural gene flow barriers to a mobile carnivore across a metropolitan area. Mol Ecol. 2020 Feb;29(3):466–84. doi:10.1111/mec.15345

56. Evans MJ, Rittenhouse TAG, Hawley JE, Rego PW, Eggert LS. Spatial genetic patterns indicate mechanism and consequences of large carnivore cohabitation within development. Ecol Evol. 2018 May;8(10):4815–29. doi:10.1002/ece3.4033

57. Anaya-Padrón MG, López González CA, Rico Y, Espinosa-Flores ME. Do highways influence the genetic structure of coyotes (Canis latrans) in a highly fragmented urban–rural landscape in central Mexico? Mammal Res. 2023 Jul;68(3):397–408. doi:10.1007/s13364-023-00692-4

58. Stevens TK, Biffi D, Chipps AS, Hale AM, Williams DA. Gene flow of small mammals is inhibited by highways and the urbanized habitat matrix in a large urban forest fragment. Ecosphere. 2025 Aug;16(8):e70343. doi:10.1002/ecs2.70343

59. Leyk S, Uhl JH. HISDAC-US, historical settlement data compilation for the conterminous United States over 200 years. Sci Data. 2018 Sep 4;5(1):180175. doi:10.1038/sdata.2018.175

60. Riley SPD, Pollinger JP, Sauvajot RM, York EC, Bromley C, Fuller TK, et al. FAST-TRACK: A southern California freeway is a physical and social barrier to gene flow in carnivores. Mol Ecol. 2006 Jun;15(7):1733–41. doi:10.1111/j.1365-294X.2006.02907.x

61. Mihalik B, Ágh N, Pipoly I, Nemesházi E, Szabó K, Seress G, et al. Low genetic differentiation and symmetric migration between urban and forest populations of great tits. Biol Futura. 2025 Sep;76(3):371–81. doi:10.1007/s42977-025-00259-1

62. Richardson JL, Michaelides S, Combs M, Djan M, Bisch L, Barrett K, et al. Dispersal ability predicts spatial genetic structure in native mammals persisting across an urbanization gradient. Evol Appl. 2021 Jan;14(1):163–77. doi:10.1111/eva.13133

63. Carlen E, Munshi-South J. Widespread genetic connectivity of feral pigeons across the Northeastern megacity. Evol Appl. 2021 Jan;14(1):150–62. doi:10.1111/eva.12972

64. Kyriazis CC, Serieys LEK, Bishop JM, Drouilly M, Viljoen S, Wayne RK, et al. The influence of gene flow on population viability in an isolated urban caracal population. Mol Ecol. 2024 May;33(9):e17346. doi:10.1111/mec.17346

65. McInturff A, Volski L, Callahan MM, Sneegas G, Pellow DN. Pathways between people, wildlife and environmental justice in cities. People Nat. 2025 Mar;7(3):575–95. doi:10.1002/pan3.10793

66. Spear JE, Grijalva EK, Michaels JS, Parker SS. Ecological spillover dynamics of organisms from urban to natural landscapes. J Urban Ecol. 2018 Jan 1;4(1). doi:10.1093/jue/juy008

67. Mannan RW, Steidl RJ, Boal CW. Identifying habitat sinks: a case study of Cooper’s hawks in an urban environment. Urban Ecosyst. 2008 Jun;11(2):141–8. doi:10.1007/s11252-008-0056-9

68. Runge JP, Runge MC, Nichols JD. The Role of Local Populations within a Landscape Context: Defining and Classifying Sources and Sinks. Am Nat. 2006 Jun;167(6):925–38. doi:10.1086/503531

69. Kanda LL, Fuller TK, Sievert PR, Kellogg RL. Seasonal source–sink dynamics at the edge of a species’ range. Ecology. 2009 Jun;90(6):1574–85. doi:10.1890/08-1263.1

70. Stillfried M, Fickel J, Börner K, Wittstatt U, Heddergott M, Ortmann S, et al. Do cities represent sources, sinks or isolated islands for urban wild boar population structure? Frair J, editor. J Appl Ecol. 2017 Feb;54(1):272–81. doi:10.1111/1365-2664.12756

71. Nisi AC, Benson JF, King R, Wilmers CC. Habitat fragmentation reduces survival and drives source–sink dynamics for a large carnivore. Ecol Appl. 2023 Jun;33(4):e2822. doi:10.1002/eap.2822

72. Gustafson KD, Gagne RB, Vickers TW, Riley SPD, Wilmers CC, Bleich VC, et al. Genetic source–sink dynamics among naturally structured and anthropogenically fragmented puma populations. Conserv Genet. 2019 Apr;20(2):215–27. doi:10.1007/s10592-018-1125-0

73. Jon P. Beckmann, Carl W. Lackey. Carnivores, urban landscapes, and longitudinal studies:: a case history of black bears. Hum-Wildl Confl. 2008 Fall;2(2):168–74.

74. Rochette NC, Rivera-Colón AG, Catchen JM. Stacks 2: Analytical methods for paired-end sequencing improve RADseq-based population genomics. Mol Ecol. 2019 Nov;28(21):4737–54. doi:10.1111/mec.15253

75. Lindblad-Toh K, Wade CM, Mikkelsen TS, Karlsson EK, Jaffe DB, Kamal M, et al. Genome sequence, comparative analysis and haplotype structure of the domestic dog. Nature. 2005 Dec;438(7069):803–19. doi:10.1038/nature04338

76. Li G, Davis BW, Raudsepp T, Pearks Wilkerson AJ, Mason VC, Ferguson-Smith M, et al. Comparative analysis of mammalian Y chromosomes illuminates ancestral structure and lineage-specific evolution. Genome Res. 2013 Sep;23(9):1486–95. doi:10.1101/gr.154286.112

77. Li H. Aligning sequence reads, clone sequences and assembly contigs with BWA-MEM [Internet]. 2013. Available from: arXiv:1303.3997v2

78. Danecek P, Auton A, Abecasis G, Albers CA, Banks E, DePristo MA, et al. The variant call format and VCFtools. Bioinformatics. 2011 Aug 1;27(15):2156–8. doi:10.1093/bioinformatics/btr330

79. Chang CC, Chow CC, Tellier LC, Vattikuti S, Purcell SM, Lee JJ. Second-generation PLINK: rising to the challenge of larger and richer datasets. Gigascience. 2015 Dec 1;4(1):s13742-015-0047–8. doi:10.1186/s13742-015-0047-8

80. Jakobsson M, Rosenberg NA. CLUMPP: a cluster matching and permutation program for dealing with label switching and multimodality in analysis of population structure. Bioinformatics. 2007 Jul 15;23(14):1801–6. doi:10.1093/bioinformatics/btm233

81. Kamvar ZN, Tabima JF, Grünwald NJ. *Poppr* : an R package for genetic analysis of populations with clonal, partially clonal, and/or sexual reproduction. PeerJ. 2014 Mar 4;2:e281. doi:10.7717/peerj.281

82. Goudet J. HIERFSTAT, a package for R to compute and test hierarchical *F* -statistics. Mol Ecol Notes. 2005 Mar;5(1):184–6. doi:10.1111/j.1471-8286.2004.00828.x

83. Bishop AP, Chambers EA, Wang IJ. Generating continuous maps of genetic diversity using moving windows. Methods Ecol Evol. 2023 May;14(5):1175–81. doi:10.1111/2041-210x.14090

84. Poisson MKP, Gebresenbet F, Butler AR, Tate P, Bergeron DH, Moll RJ. The way “urbanization” is defined has strong implications for its effects on mammal abundance. Urban Ecosyst. 2024 Dec;27(6):2367–80. doi:10.1007/s11252-024-01598-y

85. Walker K. tigris: An R Package to Access and Work with Geographic Data from the US Census Bureau. R J. 2016;Vol. 8/2(December 2016):231–42.

86. Cushing L, Faust J, August LM, Cendak R, Wieland W, Alexeeff G. Racial/Ethnic Disparities in Cumulative Environmental Health Impacts in California: Evidence From a Statewide Environmental Justice Screening Tool (CalEnviroScreen 1.1). Am J Public Health. 2015 Nov;105(11):2341–8. doi:10.2105/AJPH.2015.302643

87. Chambers EA, Bishop AP, Wang IJ. Individual-based landscape genomics for conservation: An analysis pipeline. Mol Ecol Resour. 2025 Jul;25(5). doi:10.1111/1755-0998.13884

88. McRae B, Shah V, Mohapatra T. Circuitscape 4 User Guide. Nat Conserv [Internet]. 2013. Available from: http://www.circuitscape.org

89. Mussmann SM, Douglas MR, Chafin TK, Douglas ME. BA3-SNPs: Contemporary migration reconfigured in BayesAss for next-generation sequence data. Jarman S, editor. Methods Ecol Evol. 2019 Oct;10(10):1808–13. doi:10.1111/2041-210X.13252

90. Ali OA, O’Rourke SM, Amish SJ, Meek MH, Luikart G, Jeffres C, et al. RAD Capture (Rapture): Flexible and Efficient Sequence-Based Genotyping. Genetics. 2016 Feb 1;202(2):389–400. doi:10.1534/genetics.115.183665

91. Catchen J, Hohenlohe PA, Bassham S, Amores A, Cresko WA. Stacks: an analysis tool set for population genomics. Mol Ecol. 2013 Jun;22(11):3124–40. doi:10.1111/mec.12354

92. Li H, Handsaker B, Wysoker A, Fennell T, Ruan J, Homer N, et al. The Sequence Alignment/Map format and SAMtools. Bioinformatics. 2009 Aug 15;25(16):2078–9. doi:10.1093/bioinformatics/btp352

93. Danecek P, Bonfield JK, Liddle J, Marshall J, Ohan V, Pollard MO, et al. Twelve years of SAMtools and BCFtools. GigaScience. 2021 Feb 16;10(2):giab008. doi:10.1093/gigascience/giab008 PubMed PMID: 33590861; PubMed Central PMCID: PMC7931819.

94. Foulley JL, Ollivier L. Estimating allelic richness and its diversity. Livest Sci. 2006 May;101(1–3):150–8. doi:10.1016/j.livprodsci.2005.10.021

95. Wickham J, Homer C, Vogelmann J, McKerrow A, Mueler R, Herold N, et al. The multi-resolution land characteristics (MRLC) consortium–20 years of development and integration of USA national land cover data. Remote Sens. 2014;6:7424–41.

96. Cushing L, Faust J, August LM, Cendak R, Wieland W, Alexeeff G. Racial/Ethnic Disparities in Cumulative Environmental Health Impacts in California: Evidence From a Statewide Environmental Justice Screening Tool (CalEnviroScreen 1.1). Am J Public Health. 2015 Nov;105(11):2341–8. doi:10.2105/AJPH.2015.302643

97. Wang IJ. EXAMINING THE FULL EFFECTS OF LANDSCAPE HETEROGENEITY ON SPATIAL GENETIC VARIATION: A MULTIPLE MATRIX REGRESSION APPROACH FOR QUANTIFYING GEOGRAPHIC AND ECOLOGICAL ISOLATION: SPECIAL SECTION. Evolution. 2013 Dec;67(12):3403–11. doi:10.1111/evo.12134

98. Chambers EA, Bishop AP, Wang IJ. Individual-based landscape genomics for conservation: An analysis pipeline. Mol Ecol Resour. 2025 Jul;25(5). doi:10.1111/1755-0998.13884

99. McRae BH. ISOLATION BY RESISTANCE. Evolution. 2006;60(8):1551. doi:10.1554/05-321.1

100. Wilson GA, Rannala B. Bayesian Inference of Recent Migration Rates Using Multilocus Genotypes. Genetics. 2003 Mar 1;163(3):1177–91. doi:10.1093/genetics/163.3.1177

