## Supplemental Material for "Rethink the Sink: Urban cores act as leaky sinks to maintain regional gene flow in coyotes"

### Glossary

**Urban fragmentation model:** urbanization-driven habitat fragmentation results in small "islands" of suitable habitat where populations are isolated from each other.

- Leads to an increase in genetic drift within populations and a reduction in gene flow between populations, resulting in reduced genetic diversity within populations and increased genetic differentiation between populations. This can ultimately lead to a reduction in fitness from increased drift and inbreeding.

**Urban facilitation model:** urbanization creates movement corridors or removes barriers to gene flow (physical or behavioral).

- This reduces isolation between populations, thus reducing genetic drift within populations and increasing diversity by introducing novel alleles.

**Urban sink model:** demographic model where more urban areas act as sink populations, fed by nearby exurban or wildland source populations

- **Sink population:** death rate is higher than birth rate, and population sustainability relies on immigrants from nearby source populations
  - **Attractive sink:** poor quality habitat that attracts individuals (causing negative population growth)
  - **Ecological trap:** poor-quality habitat that individuals actively prefer to settle in because it falsely signals safety or high quality
- **Source population:** birth rate is higher than death rate creating a surplus of individuals who emigrate

**Isolation by distance:** genetic distance increases with geographic distance

- This is often considered a null model, where the landscape is assumed to be fairly homogenous and distance is the only factor limiting dispersal.

**Isolation by environment:** genetic similarity is driven by environmental similarity,

- This pattern can be driven by local adaptation or very strong natal-habitat biased dispersal.

**Isolation by resistance:** treats the landscape as a heterogeneous matrix that impedes gene flow, considering not only the location of sampled individuals but the resistance values in the landscape between them.

### Supplemental Results

*Sample demographics*

We performed RADSeq on ear tissues from 260 individuals. After removing samples with insufficient coverage and filtering for unlinked, neutral SNPs, we retained 255 coyote samples with adequate data across 41,024 loci. To optimize spatial sampling density within our study area, we then selected a final subset of 220 samples located within the bounding box 33.40° N, 118.54° W to 34.28° N, 117.60° W. Of these, 107 were female, including 89 adults and 18 juveniles (subadults and pups) and 116 were male, including 97 adults and 19 juveniles (File S1). A more strictly filtered dataset was generated to calculate relatedness, resulting in a final set of 1,464 loci. Pairwise relatedness was then used to target and remove individual coyotes most related to other individuals to curate a set of unrelated samples consisting of 120 individuals (File S2). This subset of coyotes was used to validate population assignments, ensuring results were not driven only by distributions of family groups.

*Family groups (K=5) population genetics*

We identified five distinct genetic clusters (K = 5) as the optimal representation of population structure (Fig. 2), with 92 individuals (41.82%) classified as admixed (defined as <70% ancestry assigned to any single cluster). Geographically, these clusters highlight a complex interface between urban cores and exurban greenspaces. The Central Basin population was the most widespread, spanning from the southern study area to downtown Los Angeles, while the Santa Ana Mountains and Los Angeles populations exhibited high spatial breadth along coastal-to-mountain gradients. In contrast, the Palos Verdes and Crystal Cove populations were more geographically restricted. Admixed individuals were ubiquitous, occurring in all populations but most concentrated in transition zones and at the periphery of our sampling range.

Genetic differentiation among these clusters was moderate (Fst range: 0.047–0.152; Table 1). Principal coordinate analysis (PCA) confirmed this clustering, with admixed individuals occupying the intermediate space between groups (Figure 3; Figure SX). The first two axes explained 19.33% and 15.83% of the total genetic variance, respectively. The Santa Ana Mountains and Los Angeles populations exhibited the highest levels of private alleles, and Palos Verdes and Crystal Cove the lowest.

Diversity metrics revealed consistent heterozygote deficits (He > Ho) across all populations (Table 1). This deficit was statistically significant in four of the five populations, with the most pronounced effects observed in the Crystal Cove and Central Basin populations. Consequently, inbreeding levels were highest in these two populations. Notably, we identified thirty individuals (13.6%) with Fis values >=0.15 and five of those had values >=0.25, four of which belonged to the Central Basin population (File S1). Finally, allelic richness and private allele counts further distinguished these groups: the Santa Ana Mountains and Los Angeles populations exhibited the highest private allele counts, suggesting greater long-term divergence, whereas the Crystal Cove and Palos Verdes populations displayed the lowest allelic richness.

**Supplementary Figures**

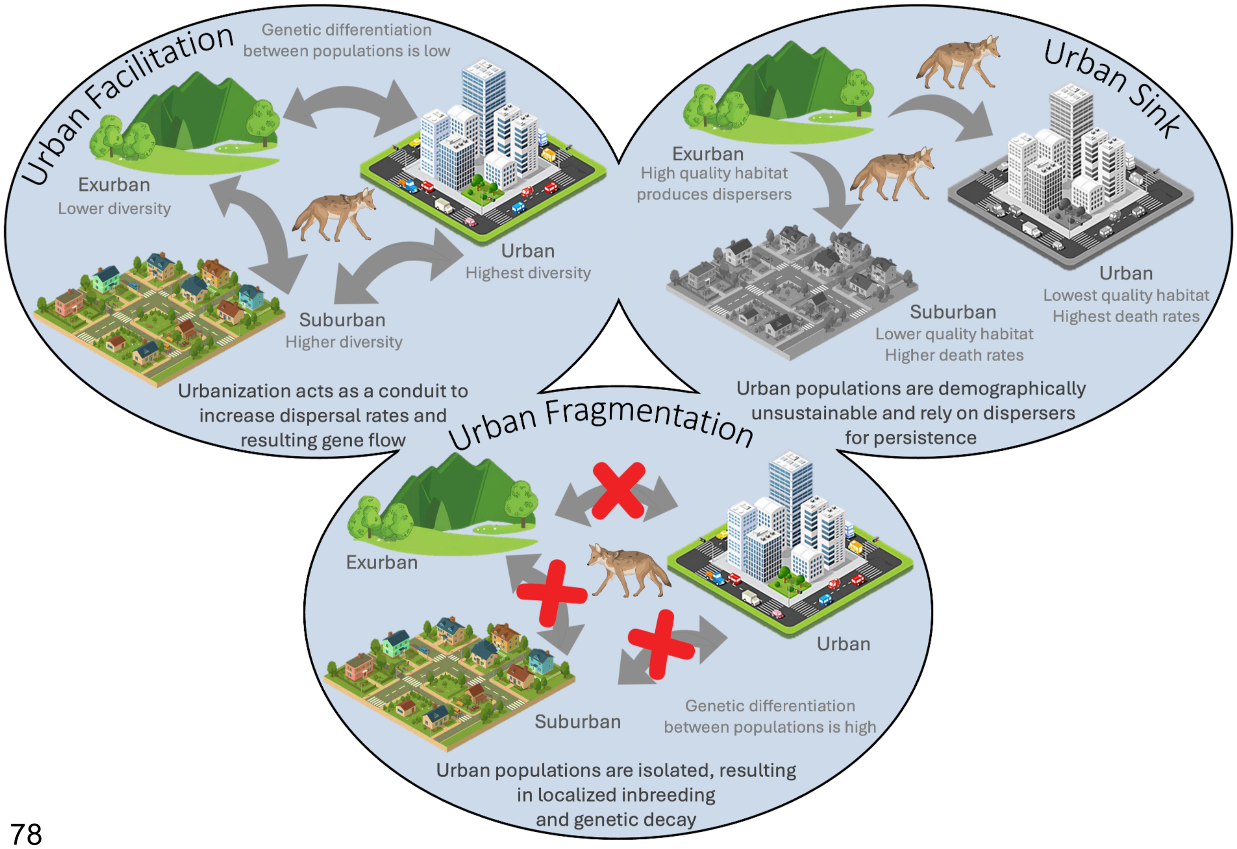


**Fig S1. Urban evolution and demographic models.** The three models tested here: Urban Facilitation, Urban Fragmentation, and the Urban Sink hypothesis.

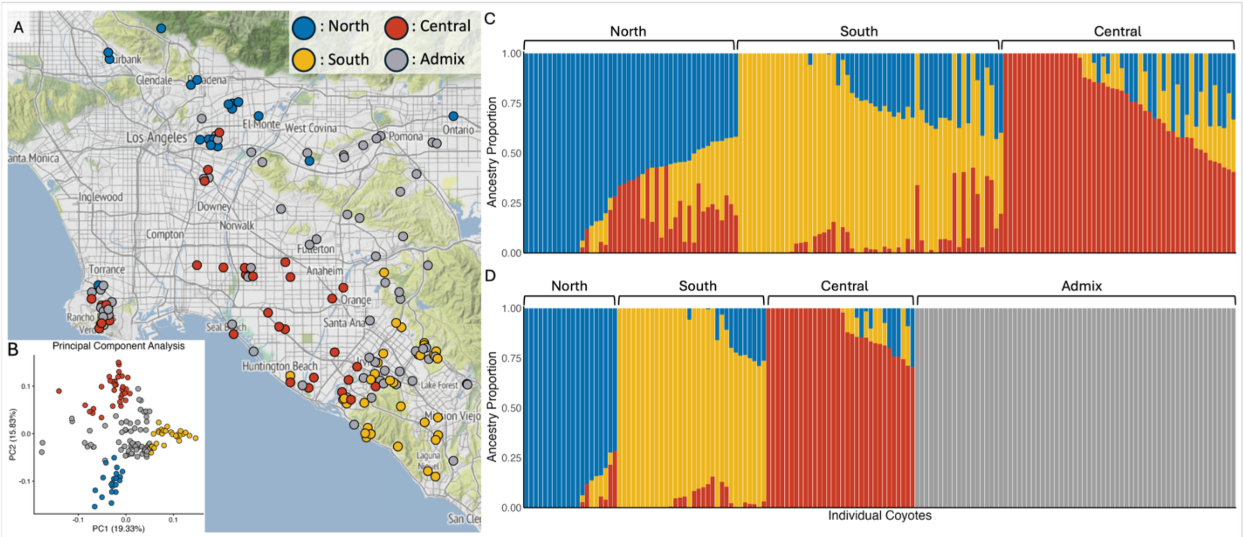


**Fig S2. Population genetic structure and spatial admixture. (A)** Spatial distribution and individual ancestry proportions mapped across the study area with admixed individuals (<70% ancestry assigned to a cluster) grouped separately. **(B)** Principal coordinate analysis (PCoA)

showing genetic differentiation along PC1 (19.33% variance) and PC2 (15.83% variance). Individual points are color-coded by assigned genetic cluster: North (blue), Central (yellow), South (red), Admixed (grey). **(C)** Ancestry proportion bar plot from admixture analysis (K = 3), where each vertical bar represents an individual and colors correspond to cluster assignment probabilities (blue = North, yellow = Central, red = South). **(D)** Ancestry proportion bar plot from admixture analysis with admixed individuals grouped separately (blue = North, yellow = Central, red = South, grey = Admixed).

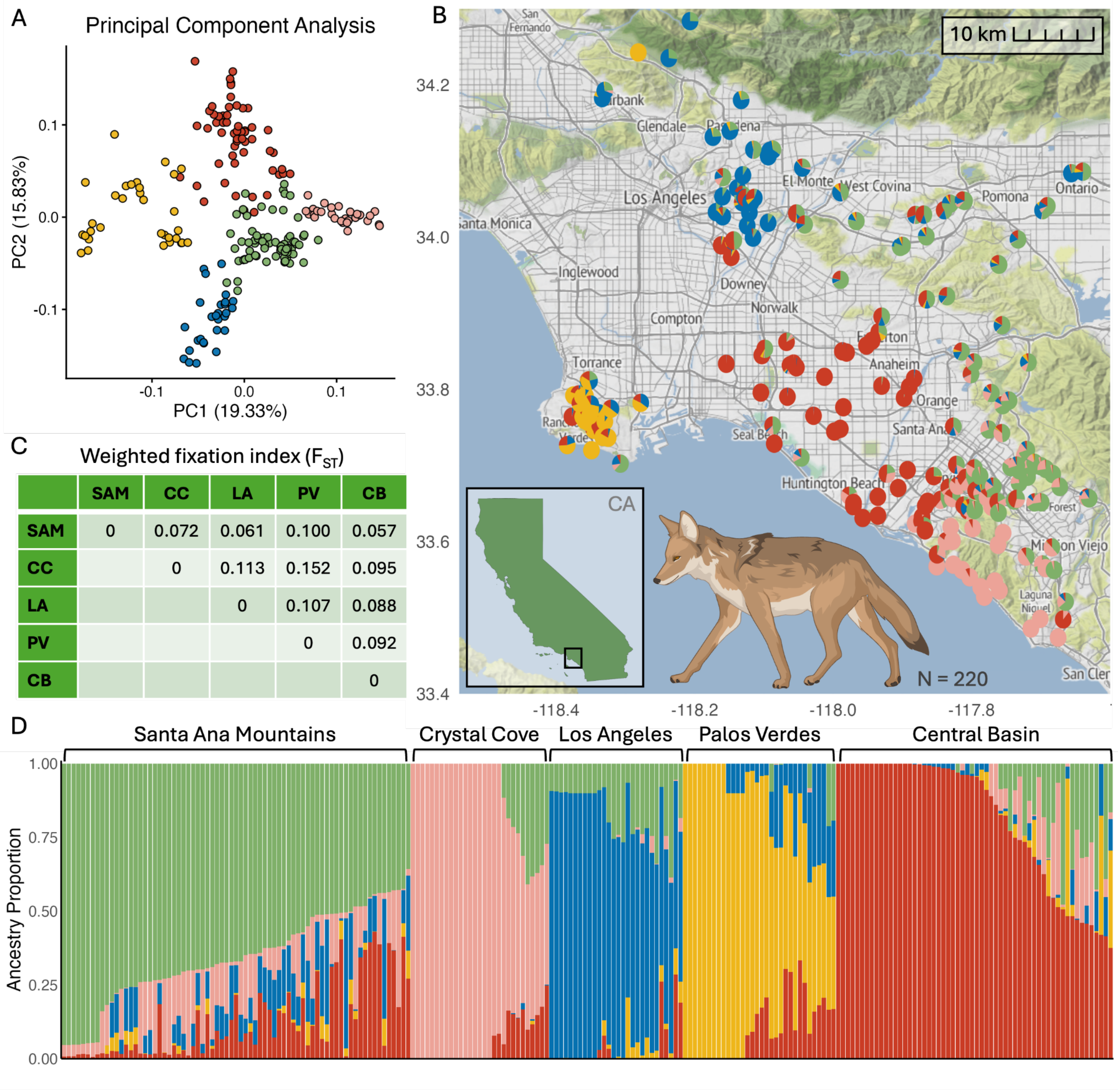


| | SAM | CC | LA | PV | CB |
|---|---|---|---|---|---|
| SAM | 0 | 0.072 | 0.061 | 0.100 | 0.057 |
| CC | | 0 | 0.113 | 0.152 | 0.095 |
| LA | | | 0 | 0.107 | 0.088 |
| PV | | | | 0 | 0.092 |
| CB | | | | | 0 |

**Fig S3. Family group genetic structure and spatial admixture (N = 220). (A)** Principal coordinate analysis (PCoA) showing genetic differentiation along PC1 (19.33% variance) and PC2 (15.83% variance). Individual points are color-coded by assigned family group: Santa Ana Mountains (SAM; green), Crystal Cove (CC; pink), Los Angeles (LA; blue), Palos Verdes (PV; yellow), and Central Basin (CB; red). **(B)** Spatial distribution and individual ancestry proportions mapped across the study area. Pie charts represent sampling locations and individual ancestry coefficients derived from K = 5 clustering with all individuals (K = 220). Inset map indicates the regional study site within California. **(C)** Pairwise $F_{ST}$ matrix showing weak genetic differentiation among genetic clusters and the admixed group. **(D)** Ancestry proportion bar plot from admixture analysis (K = 5, N = 220), where each vertical bar represents an individual and colors correspond to cluster assignment probabilities.

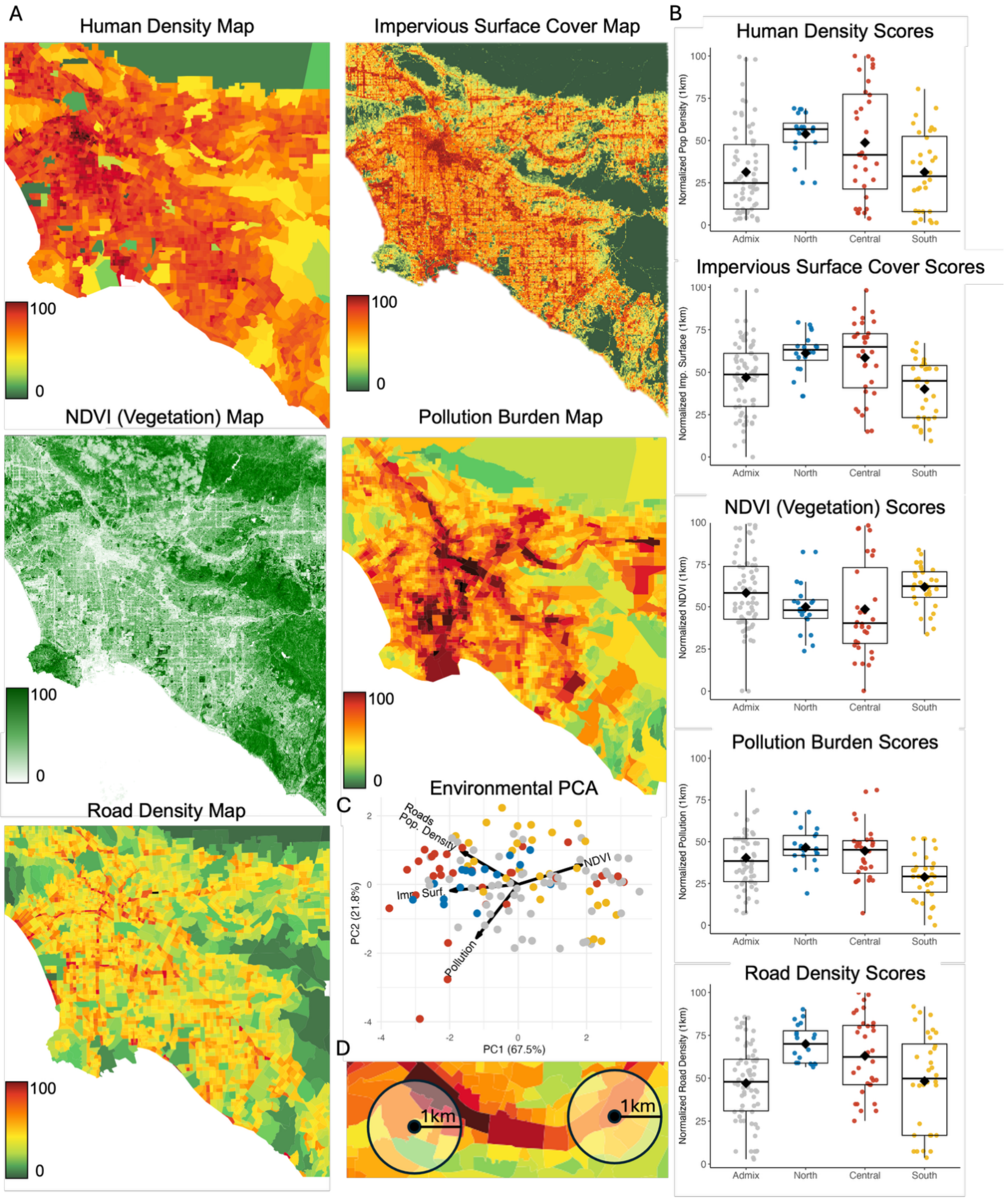


**Figure S4. Urban environmental exposure (K = 3). (A)** Maps on the left correspond to our five tested urban environmental variables. From top left to bottom: Human density, impervious surface cover, NDVI (vegetation index), pollution burden (CalEnviroScreen), and road density. **(B)** Boxplots down the right side show mean exposure across each genetic cluster (K =3 plus Admixed group). **(C)** The environmental principal coordinate analysis (PCoA) shows environmental exposure differentiation along PC1 (67.5% variance) and PC2 (21.8% variance), with arrows indicating the direction of each variable and dots colored by genetic cluster: blue = North, yellow = Central, red = South, grey = Admixed. **(D)** A schematic of the buffering around each GPS location to simulate home range.

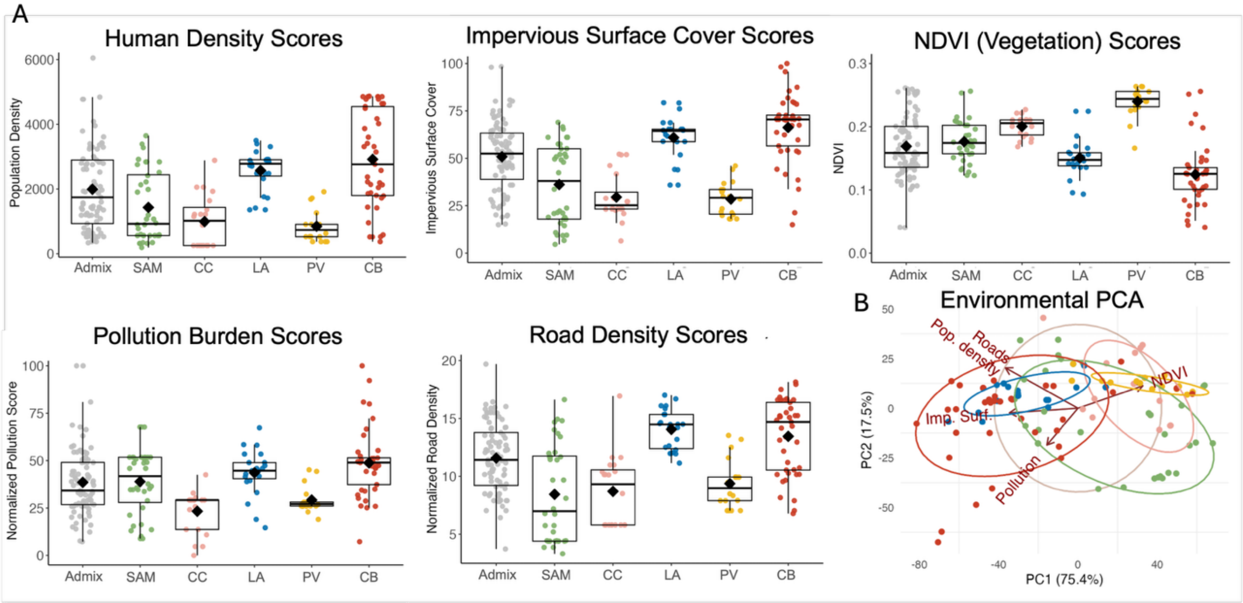


**Figure S5. Urban environmental exposure by family group (K = 5). (A)** Boxplots show mean exposure across each family group (K =3 plus Admixed group; N = 220). **(B)** The environmental principal coordinate analysis (PCoA) shows environmental exposure differentiation along PC1 (75.4% variance) and PC2 (17.5% variance), with arrows indicating the direction of each variable and dots colored by family group: Santa Ana Mountains (SAM; green), Crystal Cove (CC; pink), Los Angeles (LA; blue), Palos Verdes (PV; yellow), and Central Basin (CB; red).

*Fig S6. Isolation by Distance*

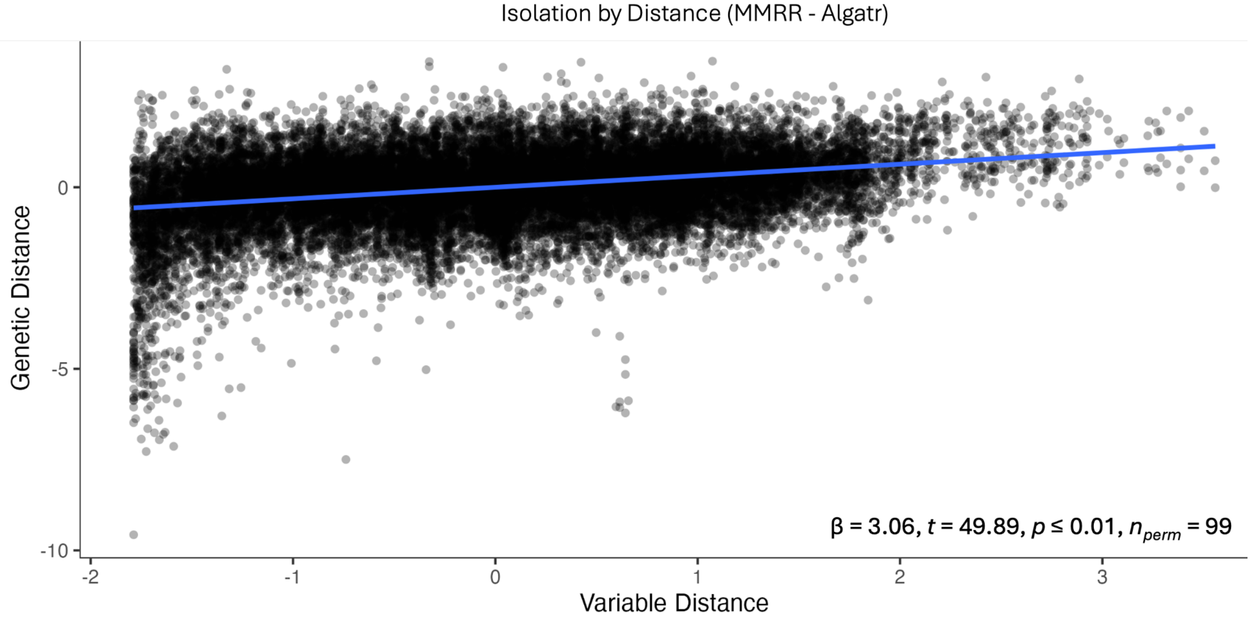


*Fig S7. BA3 Source-sink model population environmental exposure*

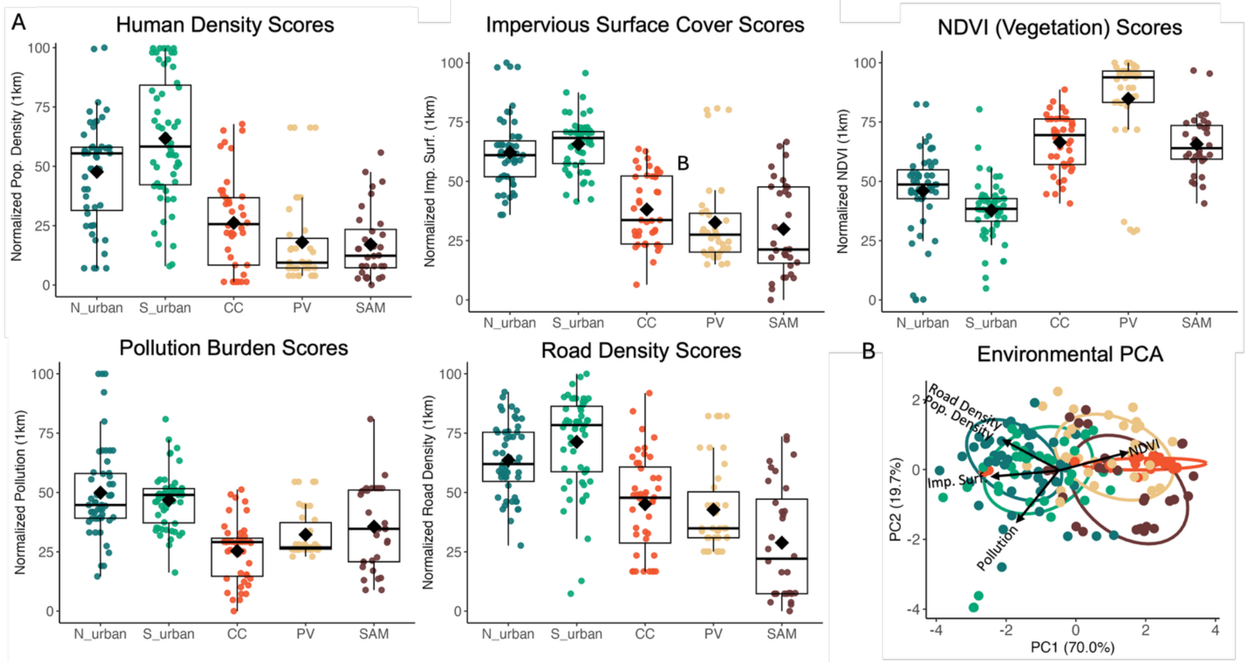


**Figure S7. Urban environmental exposure by BA3-SNPs source-sink model. (A)** Boxplots show mean exposure across each family group (K =3 plus Admixed group; N = 220). **(B)** The environmental principal coordinate analysis (PCoA) shows environmental exposure differentiation along PC1 (70.0% variance) and PC2 (19.7% variance), with arrows indicating the direction of each variable and dots colored by BA3-SNPs assigned group: teal = North urban, green = South urban, orange = Crystal Cove, tan = Palos Verdes, mauve = Santa Ana Mountains.

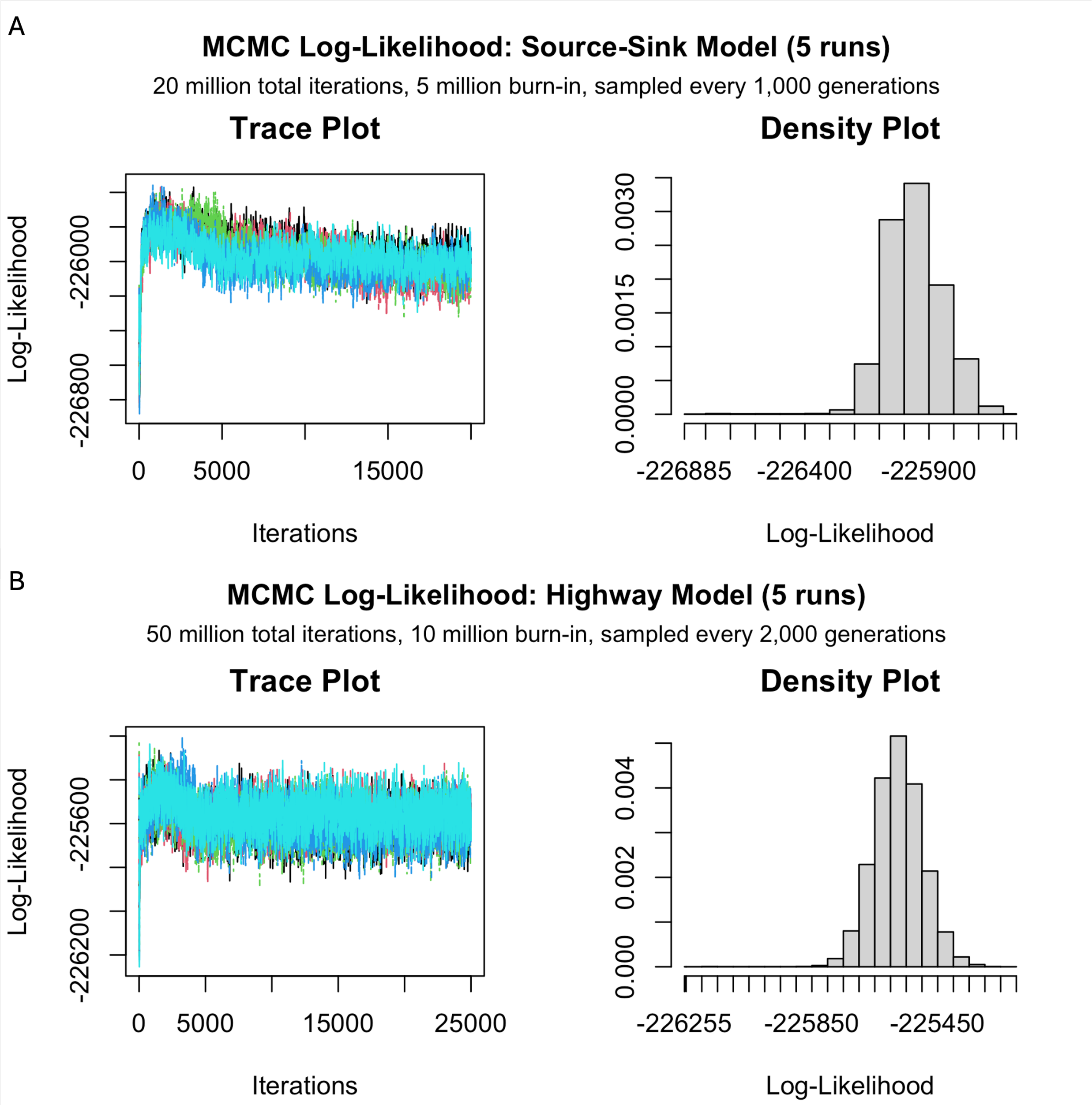


**Figure S8. Markov Chain Monte Carlo (MCMC) log-likelihood trace and posterior density plots for BA3 gene flow Source-Sink (A) and Highways (B) models.** Evaluation across five independent MCMC runs demonstrating post-burn-in convergence and stationary distributions. The left panels show trace plots of log-likelihood values across 15,000 sampled iterations (from 20 million total iterations with a 5 million burn-in, sampled every 1,000 generations). The right panels show the combined posterior probability density distribution of log-likelihood values across all chains.

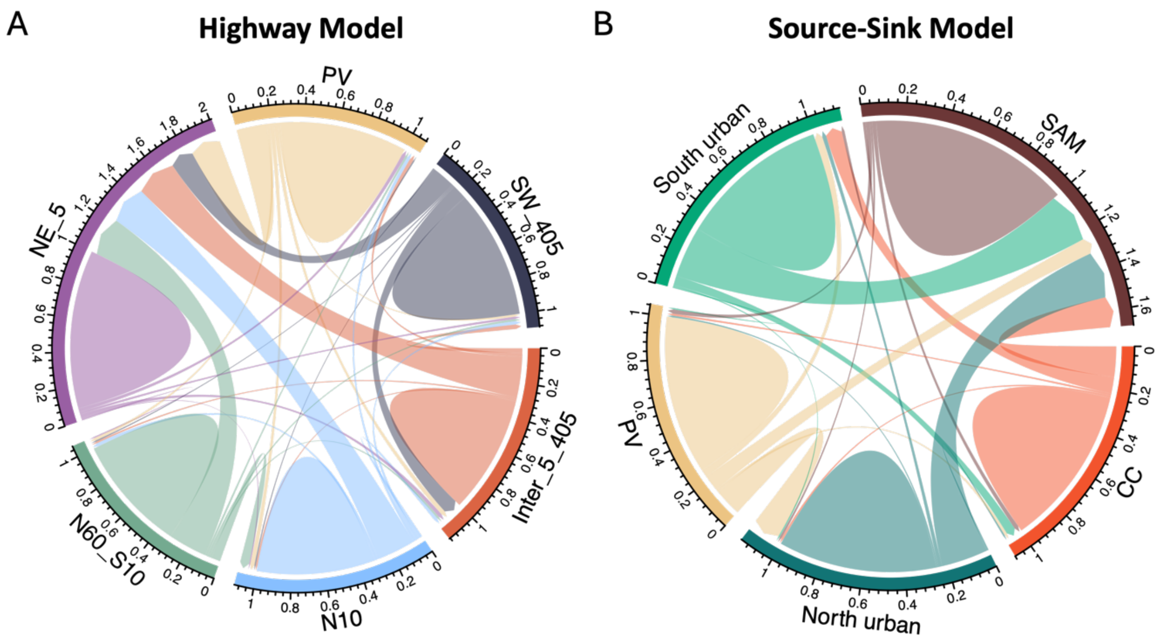


**Figure S9. Chord diagram of each BA3-SNPs gene flow Highway (A) and Source-Sink (B) models.** Ribbons indicate directional gene flow pathways, where the color and wide origin of a ribbon correspond to the source population, and the arrowhead points directly to the recipient population.

### Supplementary Tables

*Table S1. Family groups (K = 5) genetic statistics*

| Pop | N | $H_E$ | $H_O$ | t, df, p* | $F_{IS}$ | Allelic richness** | Private alleles** | Total private alleles |
|---|---|---|---|---|---|---|---|---|
| SAM | 28 | 0.207 ± 0.001 | 0.203 ± 0.017 | -1.39, 27, 0.174 | 0.022 ± 0.083 | 1.838 | 2.429 | 68 |
| CC | 20 | 0.207 ± 0.001 | 0.189 ± 0.016 | -5.19, 19, **5.24 e-5** | 0.087 ± 0.075 | 1.654 | 0.450 | 9 |
| LA | 21 | 0.207 ± 0.001 | 0.197 ± 0.016 | -2.97, 20, **0.008** | 0.051 ± 0.078 | 1.752 | 2.333 | 49 |
| PV | 18 | 0.207 ± 0.001 | 0.198 ± 0.018 | -2.31, 17, **0.033** | 0.046 ± 0.085 | 1.679 | 0.444 | 8 |
| CB | 41 | 0.206 ± 0.003 | 0.187 ± 0.02 | -6.37, 40, **1.43e-7** | 0.093 ± 0.094 | 1.725 | 0.537 | 22 |
| Admix | 92 | 0.207 ± 0.002 | 0.204 ± 0.013 | -2.05, 91, **0.044** | 0.013 ± 0.061 | 1.887 | 10.141 | 933 |

*p-values are from a Welch two-sample t-test of unequal variance between $H_E$ and $H_O$
**values are calculated on a per-individual basis to correct for uneven population sizes

*Table S2. Mean urban variable exposure score with standard deviation per ADMIXTURE population (K = 3)*

**Mean urban variable exposure score with standard deviation – no admix group**

| Pop | Human pop density | Impervious surface cover | NDVI | Pollution burden | Road density |
|---|---|---|---|---|---|
| **Central** | 43.97 +/- 32.20 | 56.76 +/- 22.39 | 51.22 +/- 27.80 | 44.38 +/- 19.98 | 59.67 +/- 21.90 |
| **North** | 38.72 +/- 23.96 | 51.98 +/- 20.24 | 57.74 +/- 21.68 | 41.92 +/- 17.06 | 55.45 +/- 20.22 |
| **South** | 31.85 +/- 23.55 | 41.96 +/- 18.23 | 58.39 +/- 13.25 | 33.57 +/- 15.32 | 46.96 +/- 27.49 |

*Table S3. Mean urban variable exposure score with standard deviation per ADMIXTURE population with Admix group separate*

**Mean urban variable exposure score with standard deviation – admix separate**

| Pop | Human pop density | Impervious surface cover | NDVI | Pollution burden | Road density |
|---|---|---|---|---|---|
| Admix | 31.29 +/- 25.58 | 46.98 +/- 21.40 | 58.22 +/- 21.76 | 40.33 +/- 18.95 | 47.09 +/- 22.44 |
| Central | 48.74 +/- 32.93 | 58.53 +/- 22.35 | 48.58 +/- 28.42 | 44.49 +/- 18.85 | 63.05 +/- 22.62 |
| North | 53.84 +/- 13.20 | 61.21 +/- 12.38 | 49.92 +/- 15.48 | 46.52 +/- 11.52 | 69.95 +/- 11.15 |
| South | 31.24 +/- 23.40 | 40.06 +/- 17.58 | 61.74 +/- 12.43 | 28.88 +/- 13.77 | 48.27 +/- 27.96 |

*Table S4. Coefficient of variation[1] (CV) of urban variable exposure with admixed separated*

Coefficient of variation[1] (CV) of urban variable exposure

| Pop | Human pop density | Impervious surface cover | NDVI | Pollution burden | Road density | Mean CV per pop |
|---|---|---|---|---|---|---|
| Admix | 81.756 | 45.554 | 37.38 | 46.984 | 47.649 | 51.865 |
| Central | 67.551 | 38.19 | 58.509 | 42.361 | 35.881 | 48.498 |
| North | 24.515 | 20.229 | 31.005 | 24.756 | 15.94 | 23.289 |
| South | 74.909 | 43.887 | 20.125 | 47.692 | 57.927 | 48.908 |
| Mean CV per variable | 62.183 | 36.965 | 36.755 | 40.449 | 39.349 | |

*Table S5. Urban covariates Pearson's correlation tests*

| | **Human pop density** | **Impervious surface cover** | **NDVI** | **Pollution burden** | **Road density** |
|---|---|---|---|---|---|
| **Human pop density** | 1 | 0.6978654 | -0.6021018 | 0.2540532 | 0.8365220* |
| **Impervious surface cover** | | 1 | -0.8794731* | 0.6023226 | 0.7597495* |
| **NDVI** | | | 1 | -0.6435622 | -0.6109679 |
| **Pollution burden** | | | | 1 | 0.2245541 |
| **Road density** | | | | | 1 |

*Table S6. Urban covariates vs genetics spearman correlation*

**Spearman correlation tests – Ho and Fis vs urban covariates (individual)**

| **Covariate** | **Shapiro-Wilk normality test (W, p)** | **$H_O$ (rho, p)** | **$F_{IS}$ (rho, p)** |
|---|---|---|---|
| Human pop density | 0.886, <0.001* | 0.107, 0.114 | -0.093, 0.172 |
| Impervious surface cover | 0.978, 0.002* | 0.049, 0.466 | -0.051, 0.456 |
| NDVI | 0.984, 0.012* | -0.064, 0.341 | 0.053, 0.436 |

| | | | |
|---|---|---|---|
| Pollution burden | 0.963, < 0.001* | 0.082, 0.226 | -0.088, 0.193 |
| Road density | 0.927, < 0.001* | -0.009, 0.892 | 0.013, 0.844 |

*Table S7. IBE GDM – Human population density*

| Predictor | Coefficient |
|---|---|
| Geographic distance | 0.74 |
| Human population density | 0.00 |
| **Deviance explained:** | **11.62%[1]** |

[1]The percentage of null deviance explained by the fitted GDM model

*Table S8. IBE GDM varimp – Human population density (1,000 permutations)*

| Predictor | Predictor importance | Predictor p-value | Model convergence |
|---|---|---|---|
| Geographic distance | 99.36 | 0.00 | 1,000 |
| Human population density | 0.92 | 0.85 | 1,000 |
| **Model deviance:** | **346.80** | | |
| **% deviance explained:** | **11.62%** | | |
| **Model p-value:** | **0.00** | | |
| **Fitted permutations:** | **993.00** | | |

*Table S9. IBE GDM – Impervious surface cover*

| Predictor | Coefficient |
|---|---|
| Geographic distance | 0.73 |
| Impervious surface cover | 0.12 |
| **% deviance explained:** | **12.23%[1]** |

[1]The percentage of null deviance explained by the fitted GDM model

*Table S10. IBE GDM varimp – Impervious surface cover (1,000 permutations)*

| Predictor | Predictor importance | Predictor p-value | Model convergence |
|---|---|---|---|
| Geographic distance | 91.27 | 0.00 | 1,000 |
| Impervious surface cover | 4.24 | 0.31 | 1,000 |
| **Model deviance:** | **344.44** | | |
| **% deviance explained:** | **12.22%** | | |
| **Model p-value:** | **0.00** | | |
| **Fitted permutations:** | **993.00** | | |

*Table S11. IBE GDM – NDVI*

| Predictor | Coefficient |
|---|---|

| | |
|---|---|
| Geographic distance | 0.74 |
| NDVI | 0.00 |
| **% deviance explained:** | **11.62%[1]** |

[1]The percentage of null deviance explained by the fitted GDM model

*Table S12. IBE GDM – Pollution burden*

| Predictor | Coefficient |
|---|---|
| Geographic distance | 0.71 |
| Pollution burden | 0.09 |
| **% explained:** | **11.93%[1]** |

[1]The percentage of null deviance explained by the fitted GDM model

*Table S13. IBE GDM varimp – Pollution burden (1,000 permutations)*

| Predictor | Predictor importance | Predictor p-value | Model convergence |
|---|---|---|---|
| Geographic distance | 92.75 | 0.00 | 1,000 |
| Pollution burden | 2.29 | 0.48 | 1,000 |
| **Model deviance:** | **345.59** | | |
| **% deviance explained:** | **11.93%** | | |
| **Model p-value:** | **0.00** | | |
| **Fitted permutations:** | **988.00** | | |

*Table S14. IBE GDM – Human population density*

| Predictor | Coefficient |
|---|---|
| Geographic distance | 0.74 |
| Human population density | 0.00 |
| **% deviance explained:** | **11.62%[1]** |

[1]The percentage of null deviance explained by the fitted GDM model

*Table S15. IBE GDM varimp – Human population density (1,000 permutations)*

| Predictor | Predictor importance | Predictor p-value | Model convergence |
|---|---|---|---|
| Geographic distance | 99.36 | 0.00 | 1,000 |
| Human population density | 0.92 | 0.85 | 1,000 |
| **Model deviance:** | **346.80** | | |
| **% deviance explained:** | **11.62%** | | |
| **Model p-value:** | **0.00** | | |
| **Fitted permutations:** | **993.00** | | |

*Table S16. IBE GDM – Road density and pollution*

| Predictor | Coefficient |
|---|---|
| Geographic distance | 0.69 |
| Road density | 0.12 |
| Pollution | 0.09 |
| **% deviance explained:** | **12.89%**[1] |

[1]The percentage of null deviance explained by the fitted GDM model

*Table S17. IBE GDM varimp – Road density and pollution (1,000 permutations)*

| Predictor | Predictor importance | Predictor p-value | Model convergence |
|---|---|---|---|
| Geographic distance | 82.97 | 0.00 | 1,000 |
| Road density | 6.46 | 0.21 | 1,000 |
| Pollution | 2.18 | 0.47 | 1,000 |
| **Model deviance:** | **341.83** | | |
| **% deviance explained:** | **12.89%** | | |
| **Model p-value:** | **0.00** | | |
| **Fitted permutations:** | **1,000.00** | | |

*Table S18. IBE GDM – Impervious surface cover and pollution*

| Predictor | Coefficient |
|---|---|
| Geographic distance | 0.70 |
| Impervious surface cover | 0.11 |
| Pollution | 0.08 |
| **% deviance explained:** | **12.47%**[1] |

[1]The percentage of null deviance explained by the fitted GDM model

*Table S19. IBE GDM varimp – Impervious surface cover and pollution (1,000 permutations)*

| Predictor | Predictor importance | Predictor p-value | Model convergence |
|---|---|---|---|
| Geographic distance | 85.71 | 0.00 | 1,000 |
| Impervious surface cover | 1.81 | 0.52 | 1,000 |
| Pollution | 3.65 | 0.34 | 1,000 |
| **Model deviance:** | **343.48** | | |
| **% deviance explained:** | **12.47%** | | |
| **Model p-value:** | **0.00** | | |
| **Fitted permutations:** | **998.00** | | |

*Table S20. IBE GDM – Human population density and pollution*

| Predictor | Coefficient |
|---|---|
| Geographic distance | 0.71 |
| Population density | 0.09 |
| Pollution | 0.00 |
| **% deviance explained:** | **11.93%**[1] |

[1]The percentage of null deviance explained by the fitted GDM model

*Table S21. IBE GDM varimp – Human population density and pollution (1,000 permutations)*

| Predictor | Predictor importance | Predictor p-value | Model convergence |
|---|---|---|---|
| Geographic distance | 92.04 | 0.00 | 1,000 |
| Human population density | 0.86 | 0.87 | 1,000 |
| Pollution | 2.31 | 0.46 | 1,000 |
| **Model deviance:** | **345.59** | | |
| **% deviance explained:** | **11.93%** | | |
| **Model p-value:** | **0.00** | | |
| **Fitted permutations:** | **998.00** | | |

*Table S22. IBE GDM HISDAC-US (1,000 permutations)*

**GDM – historical BUI exposure score**

| Year | BUI predictor | Geographic distance | Percent explained |
|---|---|---|---|
| **1900** | 0.26 | 0.73 | 12.26% |
| **1905** | 0.20 | 0.74 | 11.93% |
| **1910** | 0.20 | 0.74 | 11.93% |
| **1915** | 0.24 | 0.73 | 12.12% |
| **1920** | 0.26 | 0.73 | 12.26% |
| **1925** | 0.25 | 0.73 | 12.48% |
| **1930** | 0.18 | 0.72 | 12.22% |
| **1935** | 0.16 | 0.72 | 12.14% |
| **1940** | 0.15 | 0.65 | 12.34% |
| **1945** | 0.16 | 0.65 | 12.33% |
| **1950** | 0.10 | 0.67 | 11.95% |
| **1955** | 0.04 | 0.70 | 11.70% |
| **1960** | 0.03 | 0.68 | 11.80% |
| **1965** | 0.06 | 0.66 | 12.13% |
| **1970** | 0.06 | 0.65 | 12.41% |
| **1975** | 0.06 | 0.67 | 12.23% |
| **1980** | 0.081 | 0.68 | 12.60% |
| **1985** | 0.07 | 0.69 | 12.46% |
| **1990** | 0.05 | 0.72 | 12.11% |
| **1995** | 0.06 | 0.72 | 12.12% |

| | | | |
|---|---|---|---|
| **2000** | 0.06 | 0.73 | 12.05% |
| **2005** | 0.07 | 0.73 | 11.94% |
| **2010** | 0.11 | 0.73 | 12.11% |
| **2015** | 0.09 | 0.73 | 12.02% |

*Table S23. IBR models*

| IBR Model | Resistance Model | Correlation (r) | P value |
|---|---|---|---|
| Roads as barriers | Low (max 10) | 0.01683432 | 0.36166383 |
| | Medium (max 100) | 0.01549799 | 0.37726227 |
| | High (max 1,000) | 0.01496264 | 0.38766123 |
| Roads as corridors | Low (max 3) | 0.01706966 | 0.35936406 |
| | Medium (max 10) | 0.01741574 | 0.36456354 |
| | High (max 30) | 0.01751417 | 0.35766423 |
| NDVI | Low (max 3) | -0.0782626 | 0.95520448 |
| | Medium (max 10) | -0.0729835 | 0.93730627 |
| | High (max 30) | -0.0711118 | 0.93310669 |

*Table S24. IBR HISDAC-US*

| Year | Resistance Model | Correlation (r) | P value |
|---|---|---|---|
| 1940 | 1 – 10 | 0.02252032 | 0.32106789 |
| 1950 | 1 – 10 | 0.01996693 | 0.34026597 |
| 1960 | 1 – 10 | 0.00740747 | 0.44315568 |
| 1970 | 1 – 10 | -0.0083165 | 0.57534247 |
| 1980 | 1 – 10 | -0.0094422 | 0.57974203 |
| 1990 | 1 – 10 | -0.007868 | 0.56284372 |
| 2000 | 1 – 10 | -0.0014985 | 0.50794921 |

**Migration rates[1] – Highway population assignments**

| **Source → Dest.[2]** | **Inter 5/405** | **N 10** | **Inter 10/60** | **NE 5** | **PV** | **SW 405** |
|---|---|---|---|---|---|---|
| **Inter 5/405** | 0.689 ± 0.0001 | 0.011 0.0001 | 0.011 ± 0.0001 | **0.240 ± 0.004** | 0.022 ± 0.0001 | 0.027 ± 0.004 |
| **N 10** | 0.020 ± 0.0001 | 0.6929 ± 0.008 | 0.0233 ± 0.008 | **0.229 ± 0.007** | 0.020 ± 0.0002 | 0.020 ± 0.0001 |
| **Inter 10/60** | 0.013 ± 0.0001 | **0.059 ± 0.006** | 0.680 ± 0.0001 | **0.228 ± 0.006** | 0.013 ± 0.0001 | 0.0128 ± 0.0001 |
| **NE 5** | 0.022 ± 0.0001 | 0.007 ± 0.0001 | 0.008 ± 0.001 | 0.927 ± 0.001 | 0.022 ± 0.0001 | 0.0144 ± 0.0001 |
| **PV** | 0.026 ± 0.0002 | 0.026 ± 0.0001 | 0.0163 ± 0.007 | **0.177 ± 0.0056** | 0.744 ± 0.0003 | 0.0127 ± 0.0001 |
| **SW 405** | **0.0877** ± **0.0027** | 0.0077 ± 0.0001 | 0.0077 ± 0.0001 | **0.1371** ± **0.0027** | 0.0077 ± 0.0001 | 0.752 ± 0.0001 |

[1]Results are written as X ± Y, where X = migration rate and Y = MCMC standard deviation across 5 runs
[2]Rows represent the destination population, columns represent the source population

*Table S26. Source-sink model gene flow*

**Migration rates[1] – Source-sink population assignments**

| **Source Dest.[2]** | **CC** | **North urban** | **PV** | **South urban** | **SAM** |
|---|---|---|---|---|---|
| **CC** | **0.755 ± 0.025** | 0.010 ± 0.009 | 0.010 ± 0.009 | **0.077 ± 0.024** | **0.143 ± 0.028** |
| **North urban** | 0.008 ± 0.008 | **0.744 ± 0.023** | 0.008 ± 0.008 | 0.019 ± 0.012 | **0.220 ± 0.027** |
| **PV** | 0.013 ± 0.013 | **0.130 ± 0.032** | **0.744 ± 0.027** | 0.036 ± 0.021 | **0.076 ± 0.028** |
| **South urban** | 0.044 ± 0.018 | 0.007 ± 0.007 | 0.007 ± 0.007 | **0.703 ± 0.018** | **0.239 ± 0.026** |
| **SAM** | 0.020 ± 0.014 | 0.010 ± 0.010 | 0.020 ± 0.014 | 0.013 ± 0.012 | **0.937 ± 0.023** |

[1]Results are written as X ± Y, where X = migration rate and Y = MCMC standard deviation across 5 runs
[2]Rows represent the destination population, columns represent the source population

### Supplemental Materials & Methods

*Sample collection*

We opportunistically collected ear tissue samples from coyote roadkill carcasses or from lethal removal efforts between the years 2016-2022 across Los Angeles, Orange, San Bernardino, Riverside, and San Diego counties in southern California. This was done in collaboration with animal control agents, transportation departments, and professional trappers (File S1).

*DNA extraction and library construction*

We followed the manufacturer's protocol to obtain purified genomic DNA using the Qiagen DNEasy Blood and Tissue kit (Qiagen), which we quantified using a Qubit fluorometer system (Thermo Fisher Scientific). We then prepared the DNA for restriction site-associated DNA-sequencing (RADseq) following Ali et al. Briefly, we digested 75ng of DNA using the Sfb1-HF restriction enzyme (New England Biolabs), to which we then ligated a unique 8bp biotinylated bestRAD adapter to each specimen. We used Agencourt AMPure XP Beads (Beckman Coulter) for all purification or size selection steps. We sonicated pools of up to 48 samples to obtain a fragment size range of 300-400bp on a LE220 Covaris at the Princeton University Lewis Sigler Genomics Core facility. We then enriched each library for adapter-ligated fragments using Streptavidin Dynabeads M-280 (Invitrogen), which were then prepared using the standard protocol for the NEBNext Ultra II DNA Library Prep Kit (New England Biolabs). We incorporated the Illumina P2 i5 adapter followed by minimal amplification to incorporate a second unique barcode for each pooled library. Finally, we sequenced the genomic libraries in the 2x150nt configuration on a NovaSeq at the Yale Center for Genome Analysis to collect ~4 million reads per sample.

*Bioinformatics Processing*

We utilized several modules in *STACKS* v2.6 (74,91) to process paired-end RADseq data. First, we used the *process_radtags* module to retain reads that carried a sequenced barcode with no more than 2bp mismatches to their expected barcode sequences and had a quality score ≥10. We then demultiplexed pools and removed duplicated reads with the *clone_filter* module before mapping reads to the Canfam3.1 reference dog genome (GCA_2285.2) (75) using *bwa-mem* v0.7.19 (77). We also included the Y chromosome (KP081776.1; (76) with the reference genome for a complete representation of all autosomes and sex chromosomes. We used samtools v0.1.18 (92) to convert to the BAM format, which we then used in *STACKS* to annotate polymorphic nucleotides using the Marukilow model in the *gstacks* and *populations* modules (--vt-alpha and --gt-alpha with $p$=0.01). These steps identified and genotyped SNP loci across all the samples sequenced.

We utilized several filtering steps to retain a dataset of high-confidence loci. First, we removed loci with >10% missing data across individuals (--max-missing 0.9), excluded singleton and private doubleton variants, and removed individuals with >20% missing genotypes using *VCFtools* v0.1.17 (78,93). We further filtered the dataset in *PLINK2* (Chang et al. 2015) to remove loci with low minor allele frequencies (MAF<0.03) and high missingness (>20%). For demographic analyses that require neutral and unlinked loci, we constructed a "demographic dataset" by excluding loci in linkage disequilibrium (LD) using a moderate genotype correlation threshold ($r^2$>0.5) in *PLINK2* (--indep-pairwise 50 5 0.2) and loci that significantly deviated from Hardy-Weinberg Equilibrium (--hwe, $p$=0.001). For calculating relatedness, we applied additional filtering to retain loci with MAF>0.4 and lower levels of missing data, permitting only 5% missing data.

*Population genetics*

We conducted an initial assessment of non-model-based clustering analysis using Principal Component Analysis (PCA) to summarize the overall genetic variation. PCA was performed on the filtered genotype data using the software *PLINK2* (79). We used *ADMIXTURE* v1.3.0 (29) to conduct model-based clustering using an unsupervised maximum-likelihood framework to assign each individual coyote proportionally to *K* putative ancestral populations. We ran partitions (K) from one to 10 with the -cv flag to quantify goodness of fit. For each K (1–10), *ADMIXTURE* was run 10 times with different random seeds. Model support was evaluated using cross-validation error, summarized as mean ± SD across replicates. We used *CLUMPP* v1.1.2 (80) to generate consensus Q-matrices across replicates. Individual population assignments were determined based on the highest ancestry proportion in the consensus Q-matrix. Individuals with no ancestry proportion exceeding 70% in any cluster were classified as admixed. Finally, we visually inspected ancestry barplots and spatial patterns of individual assignments across the landscape to ensure biological interpretability.

We repeated this analysis after removing first-degree and then again removing up to second-degree relatives to separate signatures of true subpopulations from family groups. Pairwise relatedness was calculated using the *KING* function of *PLINK2* (79) with a cutoff of kinship <=0.0884. Relatedness was calculated using our more strictly filtered set of loci (see sequencing and filtering section above). Using KING output, we then targeted and removed individuals who were most related to other individuals to minimize the number of individuals removed. *VCFtools* v0.1.16 (78) was used to calculate pairwise Weir and Cockerham weighted and mean fixation indices (Fst) to quantify population differentiation.

*Genetic diversity and inbreeding estimates*

We used *VCFtools* v0.1.16 (78) for individual-level metrics of heterozygosity (observed, Ho; expected, He), and Fis, the inbreeding coefficient. We estimated allelic richness using the R package *hierfstat* (82), which we then standardized by rarefaction to address any biases due to unequal sample sizes (94). We also identified private alleles at the population level by pooling all individuals within each *ADMIXTURE*-defined population and using the R package *poppr* (81). We summed private alleles across loci to obtain the total number of private alleles per population and then divided by number of individuals in each population to account for differences in population size.

*Urban Covariates*

We investigated urban covariates that we hypothesized could influence coyote movement, health, and/or evolutionary trajectory. Importantly, we also made sure to target variables that significantly differed across our relatively small study area. Our final set of urban covariates included: human population density, impervious surface cover, Normalized difference vegetation index (NDVI), pollution burden, and road density. Because our location data for each coyote represented their place of death, we buffered each GPS point by a 1km radius to have a more accurate home range estimation (3.14$km^2$^2; Poisson et al.). The value assigned to an individual for each urban covariate was the average value from across this 1$km^2$ buffer area, excluding ocean area.

Population density values were obtained from the 2020 Census (U.S. Census Bureau) by dividing the human population count per census tract by the land area of each respective tract. Impervious surface cover was obtained from the multi-resolution land characteristics (MRLC) consortium (95) using the Fractional Impervious Surface measurements from the Annual National Land Cover Database (NLCD), which uses Landsat 8 imagery at 30m resolution. Impervious surface is defined as any processed material or structure that generates surface runoff. NDVI was generated using Landsat 8 OLI Level 1 (C2 L1) terrain-correct imaging data at 30m resolution. We filtered the date for springtime (.filterDate('2020-02-01', '2020-3-31') to maximize greenness. Cloud coverage was minimized (<10%). We used near-infrared and red bands to calculate NDVI (Equation 1) where lower values correspond to less vegetation.

$$NDVI = \frac{NIR - Red}{NIR + Red}$$

$$NIR = Near\ Infrared\ Band$$

Equation 1. Calculating NDVI

Road density values were obtained from the 2020 Census (U.S. Census Bureau) using the R package tigris v2.1 (85) by dividing kilometers of road per census tract by land area of the respective tract. We calculated a pollution burden index using CalEnviroScreen 4.0 (96). An exposure variable takes the mean of lead, particulate matter 2.5, diesel particulate matter, toxic releases, and solid waste values. An effect variable is similarly generated with groundwater threat, hazardous waste, cleanup sites and impaired water bodies. Pollution burden is then calculated per census tract using CalEnviroScreen's recommended method (Equation 2).

$$\text{EXPOSURE} = \frac{\text{LEAD + PM2.5 + DIESEL PM + TOXIC RELEASES + SOLID WASTE}}{5}$$

$$\text{EFFECT} = \frac{\text{GROUNDWATER + HAZARDOUS WASTE + CLEANUP SITE + IMPAIRED WATER}}{4}$$

$$\textbf{Pollution Burden} = \frac{(\text{EXPOSURE} + (\text{EFFECT} * 0.5))}{(1 + 0.5)}$$

Equation 2. Calculation for pollution burden index from CalEnviroScreen data

Mean exposure and standard deviations were calculated for each urban covariate across each *ADMIXTURE*-assigned population, as well as the coefficient of variance (standard deviation divided by mean). We also performed Tukey's HSD test from ANOVA to calculate pairwise comparisons of populations across urban variables. We ran Pearson correlation tests to ensure highly correlated variables (<0.7) were not included simultaneously in landscape genetics models. For visualization, each covariate scoring scale was normalized across the study area with values from 0 to 100.

*Landscape Genetics*

To test the landscape's effect on genetic findings, we began by assessing the distribution of all environmental covariates using Shapiro-Wilk normality tests. We then employed non-parametric Spearman's rank correlation coefficients (rho) to evaluate correlations between genetic statistics (e.g. Ho, Fis) and urban variables. These tests were performed both at the population (Ho, Fis, AR, PA) and individual (Ho, Fis) level.

Next, we moved to spatially explicit analyses (Table 1). We calculated the correlation between genetic distance and straight-line geographic distance (Isolation by Distance, IBD) using a multiple matrix regression with randomization (MMRR; nPerm = 1,000). We then applied generalized dissimilarity modeling (GDM; Wang) to assess the relative influence of geographic distance and environmental heterogeneity on spatial genetic variation (Isolation by Environment, IBE). Environmental metrics included human population density, impervious surface cover, NDVI, pollution burden, and road density. To assess the contribution of each predictor we also ran permutation-based variable importance analyses (gdm.varImp), which estimate how much model fit decreases when each predictor is randomly permuted to find statistically robust drivers of genetic differentiation (nPerm = 1,000). GDMs were run for each individual urban covariate in R using the *algatr* v1.0 package (98). The dependent variable in all models was the genetic distance matrix, while the independent variables included geographic distance and the given urban covariate. Multi-covariate models were run on uncorrelated environmental variables that generated positive GDM results individually (NDVI and human population density eliminated).

These models generate the same coefficient and predictor importance values for the urban variables they test, and simply reduce the relative explanatory values of geographic distance.

We examined IBR using NDVI, with higher vegetation corresponding to lower resistance, and roads, where larger road size and higher road density correspond to higher and lower resistance (roads as barriers and roads as corridors). We did not include pollution or human population density because the data was only available at a census tract level resolution. We also did not include impervious surface cover since we were already running NDVI. For each environmental variable we calculated three resistance surfaces representing low, moderate, and high resistance values. NDVI and roads as corridors were run using resistance values 1-3 (low), 1-10 (moderate), and 1-30 (high). For roads as barriers, we opted to run resistance values of 1-10 (low), 1-100 (moderate), and 1-1,000 (high).

NDVI resistance surfaces were generated directly from the rasters used in previous analyses, but resampled to 150m resolution due to computational limitations. Rather than the census tract-level road density data used in IBE analysis, we opted to use a more precise measure of road resistance. First, we created a 150m x 150m grid across our study area and then used the R package tigris v2.1 (85) to map roads across the grid. To generate a resistance score, we calculated the length of road in each cell multiplied by road type, then added together with other roads that occupy the cell. Road type was given a value based on average number of lanes (interstate = 5, U.S. highway = 4, state highway = 3, county highway = 2, local road = 1). For each resistance raster, we eliminated ocean cells as a route of movement by identifying the shoreline using the Annual National Land Cover Database (NLCD, Landsat 8 imagery at 30m resolution, obtained via MRLC (95)) and making ocean cells categorically impossible to navigate (score = NA). We used each resistance raster as the habitat input to the pairwise modeling mode of Circuitscape (v5.15.0) with coyote points input as nodes (88,99). Circuitscape uses circuit theory to model connectivity between each pair of individual coyotes, giving them a resistance score. Pairwise resistance scores were then compared against a genetic dissimilarity matrix using a partial Mantel test to quantify IBR while correcting for the effects of IBD.

To investigate whether contemporary genetic structure reflects historical urbanization patterns, we performed time-lagged IBE and IBR analyses. Historical built-up intensity (BUI) data were acquired from HISDAC-US, a raster time series detailing human development (the sum of building areas per pixel) in the US across the 19th and 20th centuries (59). We used layers for the greater Los Angeles area from 1940-2000 in five year increments at a spatial resolution of 250m. To prevent extreme urban outliers from skewing the resistance scale and to account for a suspected biological saturation in coyote avoidance of high-density areas, BUI values were capped at the 98th percentile of the modern (2015) distribution. We conducted time-lag IBE analyses using the BUI data in five-year increments from 1900–2015. IBR models were run in ten-year increments focused from 1940-2000, with higher BUI corresponding to higher resistance. The BUI resistance surface was generated directly from the raster used in IBE analyses. We ran only the moderate resistance model, with resistance values from 1-10.

Continuous surfaces of nucleotide diversity (π) were generated using moving-window and kriging algorithms implemented in the R package *wingen* (v1.0; 79). Sample coordinates were rasterized at a 1km x 1km resolution with a 5 km boundary buffer (*buffer =5*). Nucleotide diversity was calculated across a 15 km x 15 km moving window (*wdim = 15*) and only calculated for windows with >5 samples. The resulting windowed diversity estimates were spatially interpolated across the study area using ordinary kriging at native 1 km resolution (*agg_grd = 1*). To eliminate extrapolation across unsampled regions, the kriged surface was masked using the empirical 15 x 15 sample count layer to retain only grid cells supported by N > 5 samples.

To complement our landscape genomics models, we performed a spatial pedigree analysis to visualize relatedness across the landscape. Using the filtered set of 1,464 loci described above, we identified first-degree (e.g. parent-offspring, full-sibling) and second-degree (e.g. half-sibling, avuncular) relatives. We then mapped these related pairs across the study area to identify trans-barrier movement and characterize fine-scale dispersal distances occurring within the most recent one to two generations.

*Contemporary gene flow estimations and sink-source dynamics*

BayesAss3 (100) is a Bayesian approach that leverages individual multilocus genotypes to estimate recent migration rates among populations over the past few generations. Here, we apply BA3-SNPs v3.0.4 (89), a modification of BayesAss that accepts RADSeq data. BA3-SNPs requires a priori population assignments defined based on recent geography to test recent migration rates. We ran two models to test migration rates between geographic groups of interest: a “source-sink” model testing urban versus exurban groups, and a “highway” model testing inferred barriers to migration (roadways I-5, I-10, I-405, and SR-60). Analyses were performed on the subset of 1,464 loci curated from plink relatedness analyses. We used BA3-SNPs-autotune v2.1.2 (89) to set mixing parameters (source sink: --deltaM 0.10 --deltaA 0.550 --deltaF 0.0375; highway: --deltaM 0.1563 --deltaA 0.550 --deltaF 0.050). Each population assignment strategy was run five times with a different random number seed to run independent Markov chain Monte Carlo (MCMC). Both models were run with 20,000,000 iterations and a burn-in of 5,000,000; chain convergence was visualized and assessed using R (Figure S8). Migration results were then averaged across the five replicate runs.

We did not have the appropriate sampling density to test gene flow across SR-60, SR-73, SR-91, I-605, or I-710 in our highway model, or to separate out a northern exurban group (i.e. San Gabriel Mountains) in our source-sink model.
